# Active flows drive clustering and sorting of membrane components with differential affinity to dynamic actin cytoskeleton

**DOI:** 10.1101/2025.08.29.673170

**Authors:** Abrar Bhat, Amit Das, Meenakshi Iyer, Sarayu Beri, Sankarshan Talluri, Darius Koester, Madan Rao, Satyajit Mayor

**Author notes:** These authors contributed equally to this work (Abrar Bhat, Amit Das). Department of Chemical Engineering, Stanford University, CA.

## Abstract

Several cell membrane proteins, including signalling receptors and cell adhesion receptors, bind to contractile cortical F-actin at the cytoplasmic leaflet. Interaction with contractile actomyosin and associated actin flows drives these proteins into nanoscale clusters and mesoscopic assemblies of physiological relevance. In the context of diverse actin-binding membrane proteins, we ask how a specific composition of the local cluster/assembly may be achieved. Using *in vitro* reconstitution together with kinetic Monte Carlo simulations our studies reveal that an interplay between differential actin-binding affinity, protein-protein interactions and actomyosin remodelling leads to specific molecular patterning within a cluster and sorting between clusters. We show that in the lamellipodia of spreading cells *in vivo*, actin flows also result in the sorting of high and low-affinity actin-binders. This affinity-based sorting mechanism provides an attractive means for locally controlling cell membrane composition via engagement with the active cortical meshwork beneath.

## Main

A variety of functional proteins capable of interacting directly or indirectly with the actomyosin cortex, are driven into dynamic nanoclusters by contractile actomyosin flows^1–8^ (Fig. 1a). A bioinformatic investigation of membrane proteins at the cell surface indicates a diverse array of proteins are capable of this interaction and, hence, nanoclustering. About 51% of all cellular actin-binding proteins are located at the cell membrane which represents a validated set of ∼ 8% of all peripheral plasma membrane protein and 1 % of transmembrane protein species that can directly bind to cortical F-actin. This is however an underestimate of the total number of molecules since it does not include the relative abundance nor secondary interactions these actin-binding proteins can undergo with other proteins at the plasma membrane (see Supplementary Information-I). These active nanoclusters are further organized into mesoscopic assemblies, resulting in the generation of two-dimensional ‘active emulsions’ with physiological consequences^9^. This active nanoclustering mechanism, thus potentially provides a means for a local and regulated control of plasma membrane composition^10,11^. This is functionally important since many signalling receptors (T-cell receptors^12,13^, CXCR4^14^ transient receptor potential (TRP) channels^15^, EGFR^16^, AchR^17^), cell-substrate and cell-cell adhesion proteins (Integrins^18,19^, E-cadherins^4,20^) as well as glycoproteins (such as CD44^1^ and CD36^21^), and membrane-tethered outer-leaflet molecules (such as glycolipids and GPI-anchored proteins^22,23^) are subject to this clustering mechanism.

**Figure 1.**
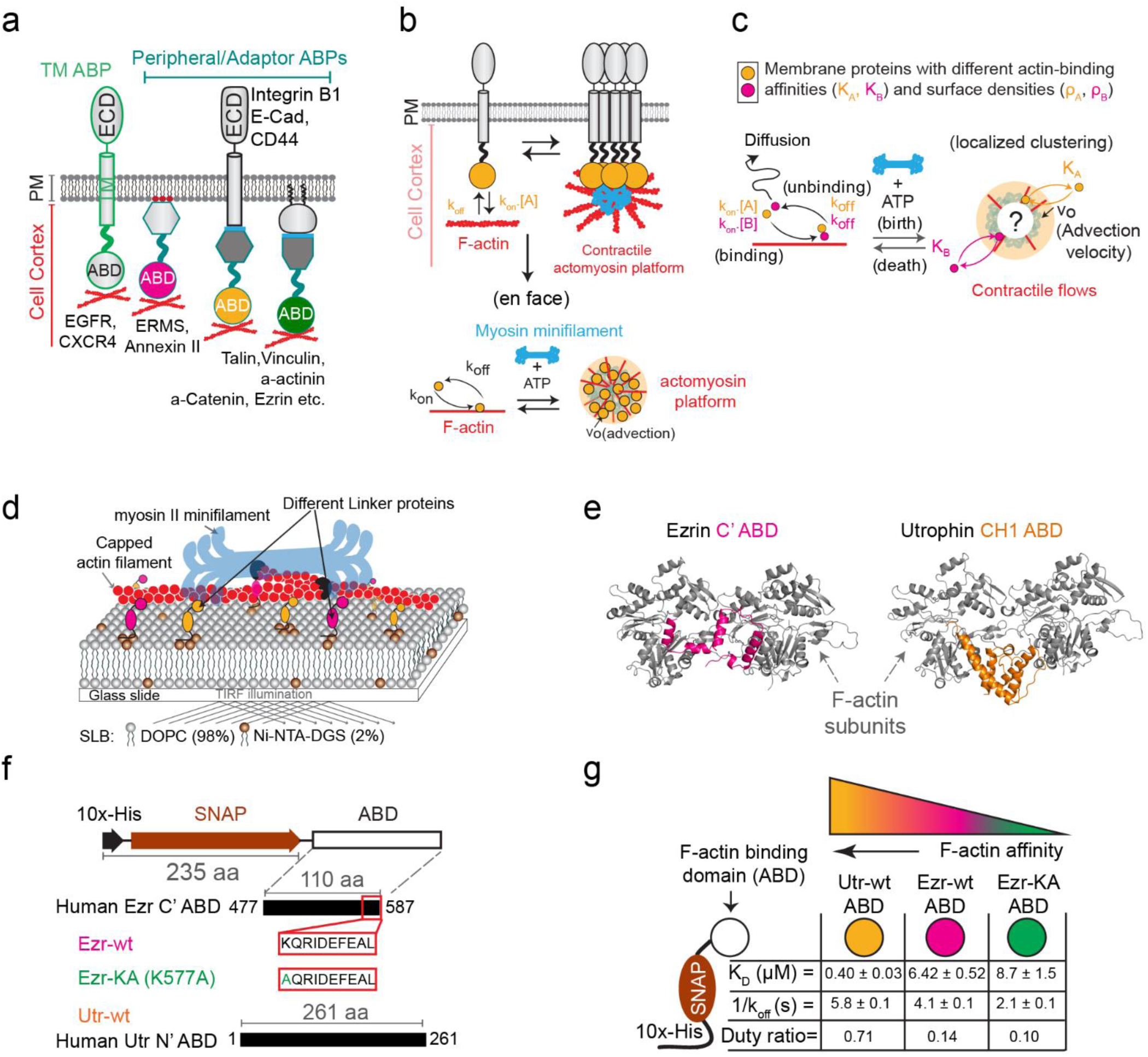
Diversity of membrane-associated actin binding proteins (ABPs) and the active clustering model. **a,** Modes of ABPs binding to cortical F-actin: either directly through actin binding domains (ABDs) of transmembrane proteins or indirectly via peripheral/adapter proteins. The diversity is indicated by the different colours of ABDs. **b**, Active clustering mechanism: a membrane-associated ABP, diffusing freely in the lipid bilayer, can be spatially clustered by contractile flows in the actomyosin cortex. The *En face* view further illustrates the active clustering mechanism. **c**, Hypothesis: differential coupling of ABPs to active cortical flows could be a mechanism for their clustering and sorting at the cell surface. **d**, In vitro reconstitution system: a glass-supported lipid bilayer with membrane-bound protein linkers capable of differential binding to self-assembling actomyosin networks juxtaposed to the bilayer. **e**, ABDs tested: the Ezrin-C’ ERM ABD and Utr-CH1-CH2 Calponin homology ABD were chosen for their significance in membrane-cortex interactions. **f**, Design of protein constructs tested: each consists of a lipid binding Deca-Histidine tag, a fluorescent tag and an ABD. **g**, Summary table of affinity measurements enumerating equilibrium dissociation constants (*K*_*D*_ = k_off_/k_on_) from F-actin co-sedimentation assay, dwell time (1/k_off_) from a single molecule TIRF assay, and duty ratio [calculated as *K*_*ABD*_ = *k*_*on*_/(*k*_*on*_ + *k*_*off*_)] (see Supplementary Fig. 2 for experimental details).

Often the efficiency of signalling depends on the ability to cluster a fixed number of a specific species of a signalling protein^24–27^. Given the diversity of actin-binding proteins (Supplementary Information-I Fig. SI1), how does this generic active mechanism give rise to the molecular specificity of the functional clusters, since all it requires is for the plasma membrane protein to acquire actin binding capacity, either directly via their cytosolic actin-binding domains (ABDs) or indirectly via actin-binding adaptors and cofactors^6–8^ (Fig. 1a, Supplementary Information-I Fig. SI1a,b)?

With limited and often overlapping interaction sites on F-actin^28^, these ABDs have diverse ways of interacting with actin^8,29^ (Supplementary Information-I Fig. SI1c) featuring a range of binding affinities for cortical F-actin^29–34^. Thus, we hypothesise that this active mechanism could give rise to a regulation of the compositional specificity of functional active nanoclusters^35–37^ based on the range of binding affinities (Fig. 1c) of the inherent diversity of membrane-associated actin-binding proteins capable of being clustered (Fig. 1a, b).

Here we demonstrate, using a theoretical description of actomyosin-induced protein clustering^5,38^,*in vitro* reconstitution of a thin layer of actomyosin on a supported membrane bilayer^39,40^, and *in vivo* experiments, how such a specificity may be achieved. In our *in vitro* system, we recreate a dynamic actomyosin network coupled to a supported lipid bilayer via membrane-actin linker proteins with varying affinities for F-actin. We demonstrate active patterning and affinity-based sorting of these linker proteins driven by contractile actomyosin flows. Our theoretical description is based on a kinetic Monte Carlo simulation^5,38^, motivated by our earlier work on active hydrodynamics of the actin-membrane composite^2,41^. We find that the interplay between differential actin-binding affinity, protein-protein interactions and actomyosin remodelling can lead to specific molecular patterning within a cluster and sorting between clusters. This type of sorting is also realized in live cells in the context of actin-driven flows during lamellipodial extension and retraction^42,43^.

## Results

To address how active contractile actin flows are generated and result in a specific lateral organization of membrane proteins with diverse F-actin-binding affinities (Fig. 1c), we simultaneously explore this in an in-vitro system40 (see also Supplementary Fig. S1d-n, Supplementary Information-II) and a kinetic Monte Carlo simulation5,38 (see Supplementary Information-III). These studies recapitulate the salient features of active cell membrane organization39,40. The simulations make further predictions that we realize in vitro and validate a key prediction in an in vivo context.

### Actin binding membrane proteins are sorted by active contractile flows

We use our established *in vitro* system^39,40^, which mimics the membrane-actomyosin composite^44–46,39^. Briefly, we hierarchically assembled it from a simple supported lipid bilayer (SLB) decorated with various membrane-actin linker proteins, added in pre-specified combinations and concentrations followed by the addition of pre-polymerized actin filaments of defined lengths, myosin II motors and ATP to generate contractile actomyosin flows that we visualize utilizing multi-colour TIRF (total internal reflection fluorescence) microscopy (Fig. 1d).

Here, we utilize three different membrane-actin linkers sharing a set of common features, a lipid-bilayer binding tag, a fluorescent marker, and an actin-binding domain (ABD) with a range of F-actin affinities spanning from ∼400 nM to ∼10 μM (Fig. 1e-g). To characterise the binding affinities of these membrane-actin linkers, a F-actin co-sedimentation assay was used to determine the equilibrium dissociation constant, *K*_*D*_ (k_off_/k_on_) and a TIRF-based single-molecule actin binding assay to measure the dwell time (1/k_off_). These assays yielded a detailed characterization of the three types of linkers used (Fig. 1g, Supplementary Fig. S2). The equilibrium affinity values were further converted into duty ratios, *K*_*ABD*_, [calculated as *K*_*ABD*_ = *k*_*on*_/(*k*_*on*_ + *k*_*off*_)]. The table (in Fig. 1g) shows the measured duty ratios of the three binders, the strongest being wild type Utrophin-ABD, Utr-wt (*K*_Utr−wt_= 0.71), followed by wild-type Ezrin, Ezr-wt (*K*_Ezr−wt_ = 0.14) and its actin-binding mutant, Ezr-KA (*K*_Ezr−KA_ = 0.10).

We first investigated the case where same amounts of two binders with different actin affinities are driven by contractile actin flows as schematized (Fig. 1c). For this, Ezr-wt and Ezr-KA were added to the SLB at equal concentrations to generate equal surface densities due to their shared SLB binding properties (10xHistidine-tag interacting with Ni^2+^-chelated lipids). Without F-actin, both the binders show free lateral diffusion (Supplementary Fig. S2o) and a uniform spatial distribution on the SLB (Supplementary Fig. S3a). After addition of F-actin, at steady state, Ezr-wt shows a robust colocalization with F-actin (Supplementary Fig. S3b) and a significant reduction in its apparent diffusion coefficient (Supplementary Fig. S2o-q). Likewise, the weaker F-actin binder Ezr-KA, shows a significantly reduced lateral mobility in the presence of F-actin, when it is not competing with Ezr-wt (Supplementary Fig. S2o-q), but is homogenously distributed in the presence of the stronger binder Ezr-wt (Supplementary Fig. S3a, b).

Upon introduction of myosin, actin filaments at first display typical active flows driven by myosin contractility, organizing into distinct and concentrated actomyosin platforms/asters that exhibit several cycles of remodelling (Fig. 2a-h; see also Supplementary Fig. S1f, I; and Supplementary Movie 1; and Refs^2,39^). At high ATP levels, actin asters display rapid remodelling, visualised as a transient local enrichment and dissipation of actin density (Fig. 2a, b). This affects the strong and weak binders differently where the stronger binder, Ezr-wt, is concentrated at the core of the asters while the weaker binder, Ezr-KA, is excluded (Fig. 2 b-e; Supplementary Fig S4a, c). Though the sorting is transient and occurs only during the local enrichment of actin in the fast-remodelling asters, an average over time provides definite evidence for micro phase segregation (∼200-450 s in Fig. 2b, 2f; see also *Methods*). When the actomyosin complexes disassemble (between 480-750 s in Fig. 2b, 2f), the local membrane composition resets to its basal state (Fig. 2f). The two binders therefore undergo an actively driven spatial sorting in the absence of any specific intermolecular interaction (Fig. 2f; see *Methods*). At low ATP levels, aster remodelling reduces gradually^48,49^, leading to a jammed state of the acto-myosin assembly and an eventual decrease in the sorting ratio, with the binders reverting to a more equilibrium distribution (Fig. 2g-k; Supplementary Fig. S4a-i; Supplementary Movie 2, see *Methods*).

**Figure 2.**
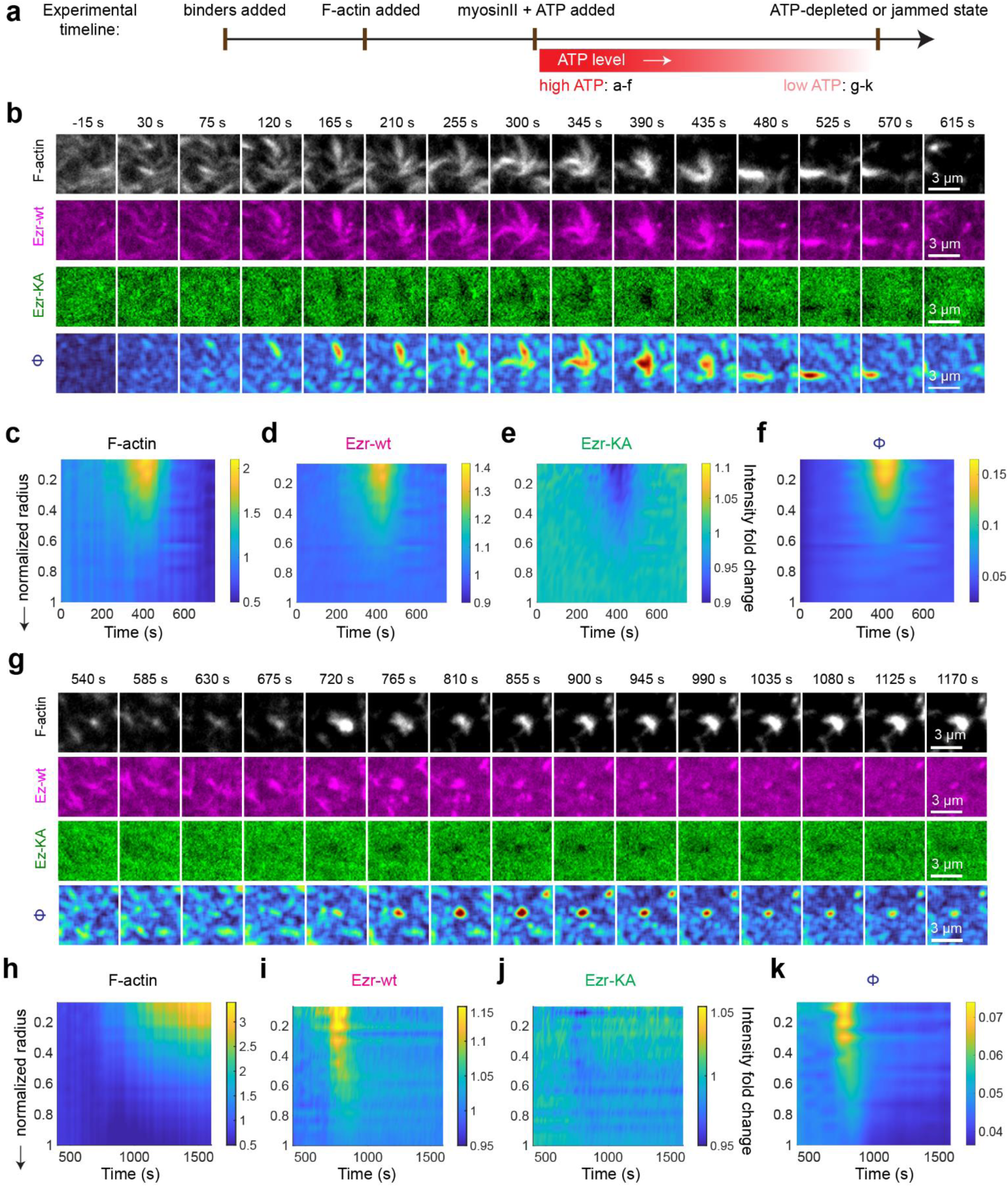
Active clustering and spatial sorting of two ABPs driven by remodelling contractile asters in supported lipid bilayers. Experimental timeline depicts the order of addition in the *in vitro* supported bilayer assay **a**, TIRFM images of F-actin (grey), Ezr-wt (magenta, stronger F-actin binder) and Ezr-KA (green, weaker F-actin binder) at different time points show transient, localized clustering of Ezr-wt and concomitant exclusion of Ezr-KA, driven by actomyosin flows (between t = 165-435 s). Myosin was added at t = 0 s. The sorting pattern dissipates as the contractile actomyosin platforms actively disassemble (480- 615 s). **b-d**, Normalized radial intensity maps of actin, Ezr-wt (magenta) and Ezr-KA (green) in high-ATP remodelling aster state (n = 21). **e**, radial map of the segregation parameter φ [= (I_A_ - I_B_)/(I_A_ + I_B_), where I = normalized intensity of respective binders; see Methods] represents the extent of spatial segregation between the binders. φ = 0 means complete mixing; φ =1 means complete segregation; 0 < φ < 1 means partial segregation. Note: a transient increase in φ at t ∼400 s coincides with the formation of localized contractile asters (peak in b) and returns to 0 as actin intensity decays due to an active disassembly of the aster (see movie 1). **f-j**, Spatial and temporal organization of adapter proteins in the low-ATP, jammed aster phase. **f**, TIRF images of the three proteins at longer time points when the ATP levels are low. **g-I**, Normalized radial intensity colour maps showing a gradual increase in actin intensity (**g**), localized clustering of Ezr-wt (**h**) and a concomitant depletion of Ezr-KA (**i**), driven by the jamming asters (n = 41). **j**, Radial colour map of φ in the jammed aster state, showing modest sorting compared to (**e**). φ resets to 0 as the asters approach the ATP-depleted jammed state (t ∼ 1250 s). Scale bar, 3 μm.

### Simulations predict steady state patterning driven by contractile flows and non-equilibrium remodelling

The observed protein sorting in the remodelling phase can be understood using a kinetic Monte-Carlo simulation^5,9^ describing the clustering of membrane proteins that interact with the active actomyosin fluid^2,41,50^. We had earlier shown^5,9,41^, and observed in experiments^2,39^, how actin flows generated within the contractile actomyosin fluid, accompanied by turnover, naturally drive the continual assembly and disassembly of finite size asters. Here we represent the transient contractile asters by a finite density of non-overlapping circular discs in two-dimensions that appear and disappear stochastically (Fig. 1c, see *Methods* and Supplementary Information-III). These discs essentially represent domains of the cell cortex where the magnitude of contractile stresses of actomyosin is high. From previous work, the distribution of remodelling timescale of these contractile domains is taken to be exponential^38,51^, representing a Poisson distributed birth-death process.

As described earlier^38^, membrane proteins that bind to actin are advected by the contractile flows and driven to cluster at the core of transient asters. In the stochastic simulation, we represent this by having membrane proteins move diffusively (with diffusion coefficient, D) when not influenced by the contractile domains, and advected with a centripetal velocity v_0_, when interacting with the contractile domain. Here, we study the dynamics of two protein species, A and B, with densities n_A_ and n_B_, respectively. Proteins A and B interact amongst themselves via a weakly attractive Lenard-Jones (LJ) potential, ∈_*AA*_= ∈_*BB*_= 0.1*k*_*B*_*T*, while the A-B interaction is taken to purely repulsive LJ. The interaction of the proteins with the contractile domains is parameterised by binding-unbinding rates (or equivalently duty ratios), corresponding to different binding affinities, K_A_ and K_B_ (Fig. 1c). For details of the simulation parameters and the precise implementation of the kinetic Monte Carlo simulation (see *Methods* and Supplementary Information-III).

Starting from a uniform distribution of A and B, we systematically vary three parameters: i) the relative affinity (ratio of duty ratios = K_A_/K_B_); ii) the relative density (n_A_/n_B_); and iii) the mean remodelling timescale *τ*_*rem*_ of the contractile domains, to create a nonequilibrium phase diagram of micro phase segregated patterns (Fig. 1c, Fig. 3a-c).

**Figure 3.**
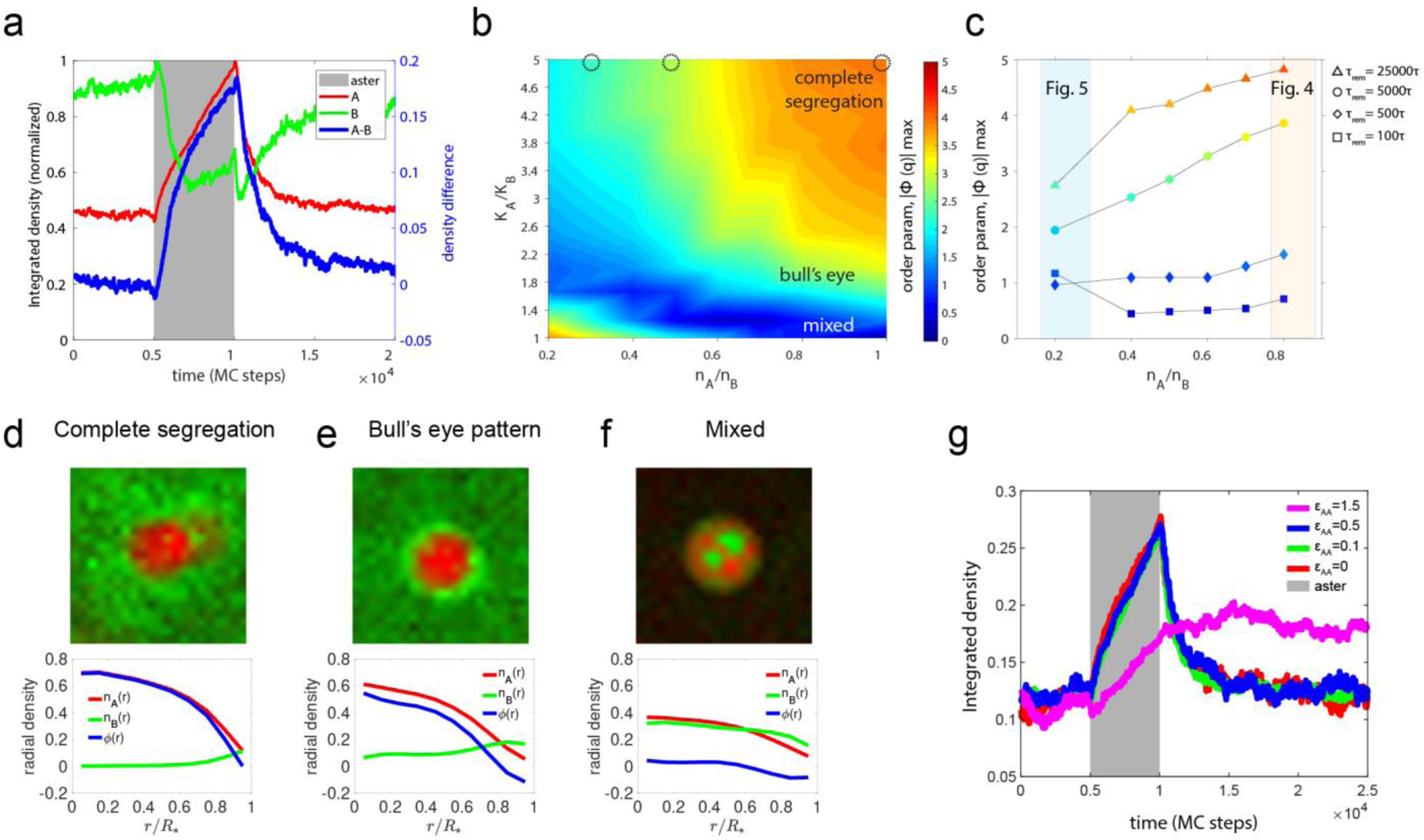
Actomyosin templated contractile platforms drive steady state sorting and patterning of membrane components with differential actin-binding affinity. **a**, Time-series of local actin concentration driven by myosin contractility, along with normalized densities of actin-binding molecules A and B and density difference (I_A_ - I_B_), exhibit sorting (blue line) as a consequence of actomyosin based contractile flows (grey zone corresponds to the aster) (SI Movie 3). All densities are normalized to the maximum. **b,** Phase diagram of steady-state patterns from the simulation at *τ*_*rem*_ = 5000 *MC steps*, mapped as a function of affinity ratio K_A_/K_B_ and the number density ratio n_A_/n_B_. We have varied K_B_ relative to a fixed K_A_ = 0.30. Heat-map shows how the segregation order-parameter (|Φ_*q*_|_*max*_) changes in this phase space. |Φ_*q*_|_*max*_is the absolute value of the strongest Fourier mode of the radial density difference *ϕ*(*r*). Three distinct micro-phase segregation patterns emerge: (i) complete segregation of A clusters (**d**), (ii) a bull’s eye pattern with A in the inner core **e**) and (ii) a mixed state (**f**). High φ values correspond to complete segregation (**d**), while φ decreases in the bull’s eye pattern (**e**), and goes to zero in the mixed phase (**f**). **c**, Effect of aster remodelling time, *τ*_*rem*_, on the active patterning of the binders. For a fixed relative affinity (K_A_/K_B_ = 5) and varying density ratios (n_A_/n_B_), patterning transitions from mixed to complete segregation as *τ*_*rem*_ is increased from 100 to 25000 MC steps. Longer *τ*_*rem*_results in slower aster turnover and longer aster lifetimes, leading to complete segregation. The colours of the symbols follow the same colour scale as in panel b to help identify which type of pattern this corresponds to. **g**, Time-series of density of stronger binder A for variable values of like-like interaction strength for the advected species, ɛ_AA_ = 0, 0.1, 0.5, 1.5 in units of k_B_T. Note clusters persist even after the contractile platform has disappeared with interactions stronger than thermal energy scale (ɛ_AA_= 1.5). For all the cases in g, we have used K_A_/K_B_ = 10.

In general, when K_A_ > K_B_, molecules of A interacting with the contractile domain get advected sooner towards the domain core, excluding the weaker binding B proteins due to steric repulsion. When the contractile pattern disassembles, the stronger binder enriched at the domain core diffuses away, locally decaying to the original mixed profile (Fig. 3a, see Supplementary Movie 3).

We first characterize the spatial patterning as a function of K_A_/K_B_ and n_A_/n_B_, keeping the remodelling rate of the contractile domains fixed, relative to the diffusion rate of the proteins over the scale of the domain. Microphase segregation is described by the local order parameter profile over the scale of the domain, *ϕ*(*r*) = 〈*n*_*A*_(*r*)〉 − 〈*n*_*B*_(*r*)〉 where *n*_*A*_(*r*) and *n*_*B*_(*r*) represent the radial density of A and B, respectively at a distance *r* from the centre of the domain. The angular brackets represent spatial and temporal averages taken over annular regions centred around the domain-core with mean radius *r* and the lifetime of the contractile domain (see Supplementary Information-II for details). We observe three distinct microphases (Fig. 3b, 3d-f): (i) a mixed pattern, with A and B co-clustered within the contractile domain, (ii) a bull’s eye pattern, with A at the domain core and B forming a ring at the periphery, and (iii) a complete segregation, with B excluded from the clusters of A.

For K_A_/K_B_ > 1, the segregation parameter decreases when the mean remodelling timescale of the domain increases (Fig. 3c), meaning that the contractile domains do not have enough time to completely advect the stronger binder towards the domain core. In earlier work^2,38,52^ we had demonstrated, how a uniform distribution of a single protein species (A) in an active fluid, segregates into domains of A-rich and A-poor regions even in the absence of an attractive interaction between the A proteins. In a similar manner, the microphase segregation observed here persists even when there is no attractive interaction between the proteins, reaffirming the nonequilibrium origins of this segregation (Fig. 3g). Consistent with this, we find that it is possible to drive the system from one pattern to another simply by tuning *τ*_*rem*_, keeping other parameters fixed. Figure 3c illustrates many ways of traversing these phases, including re-entrant behaviour. We remark that these distinct microphases show up even in the low *τ*_*rem*_(i.e. fast remodelling) regime, when we average the order parameter over many remodelling events.

### Comparison of theory with *in vitro* experiments

Our reconstitution system together with the array of actin-binding proteins that we have synthesized, allows us to experimentally scan through the phase diagram obtained from our simulations of this nonequilibrium system.

#### Effect of relative actin binding affinities and remodelling rates of the asters

To explore a representative region of the phase space indicated by the simulation (shown as dark circles in Fig. 3b), we first ascertain that our simulations recapitulate the *in vitro* results obtained with K_Ezr-wt_/K_Ezr-KA_ = 1.4 and n_Ezrwt_/n_Ezr-KA_ =1 (compare Fig. 2b-f, with Supplementary Information-III Fig. S2 and Fig. 3b) where we observe partial spatial segregation of the higher affinity binder upon aster remodelling.

We next explored an area of the phase diagram predicted by the simulations (shown as dark circles in Fig. 3b and as orange box in Fig. 3c), where we change the affinities of the binders by utilizing different actin-binding linkers with varying binding affinities to explore their behaviour in pairwise combinations. When K_Utr-wt_/K_Ezr-wt_ = 5 (at n_Utr-wt_/n_Ezr-WT_ =1) we find that the relatively stronger binder, Utr-wt (Fig. 1g), exhibits a robust, enrichment with actin (Fig. 4a, c), and the relatively weaker binder Ezr-wt, instead of being excluded, also exhibits a modest enrichment in the asters along with Utr-wt (Fig. 4a, d, see Supplementary Movie 4). This reflects a mixed phase, consistent with the region of the predicted phase space being sampled (Fig. 3b, c and Supplementary Information-III Fig. S2).

**Figure 4.**
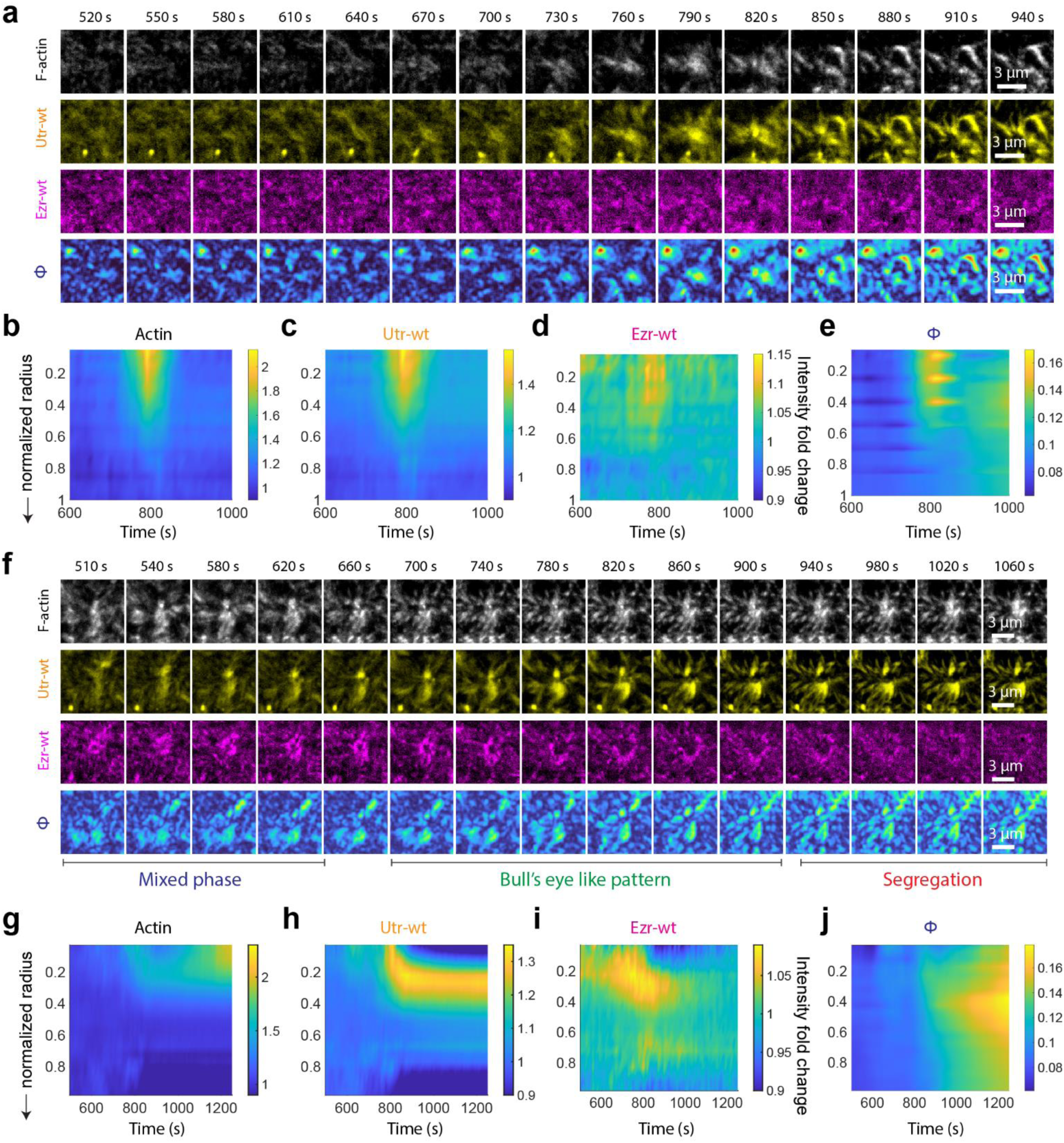
Contractile actomyosin flows on a supported bilayer recapitulate spatial patterning of membrane components with differential actin-binding affinity. (**a-e**) Spatial organization during the high-ATP remodelling aster state. TIRF images (**a**) and normalized radial intensity maps (**b-d**) show F-actin, Utr-wt (stronger F-actin binder) and Ezr-wt (weaker F-actin binder in this case) during the remodelling state. Utr-wt exhibits strong clustering whereas Ezr-wt shows only mild clustering (n = 10 asters; see Fig. S4 for details). **e,** Radial map of φ shows a mild increase at peak actin enrichment (∼800 s), despite a >1.5 peak fold change in Utr-wt, suggesting mixed co-clustering rather than exclusion. The BSR returns to 0 as the actin intensity decays (∼900 s). (**f-j**) Spatial organization during the low-ATP jammed aster state. TIRF images (**f**) and normalized radial intensity maps (**g-i**) show localized clustering of Utr-wt and a transient co-clustering of Ezr-wt driven by contractile actomyosin flows (t ∼800 s). F-actin enriches centripetally, peaking at the centre of the aster. Utr-wt begins to enrich at the centre but shifts slightly outwards at later time point (850 s onwards) and remains clustered even after asters enter the jammed state (yellow montage). Ezr-wt initially co-clusters with Utr-wt followed by a bully’s eye pattern, with Utr-wt concentrated close to the centre (peak at ∼ 0.2 normalized radius) and Ezr-wt at the periphery (peak at ∼ 0.6 normalized radius) (∼ 850-1000 s). This state is followed by a significant exclusion of Ezr-wt from Utr-wt clusters (1000 s onwards). (**j**) Radial map of φ in the jammed state shows an increase in the jammed state (∼1050 s onwards). Scale bar, 3 μm.

To examine the influence of the remodelling rate of the contractile asters on the types of patterns we explored a relationship between the remodelling rate and the pattern that develops. In addition, we also allow the *in vitro* system to deplete the levels of ATP by the continuous turnover of the energy consuming myosin motors^49^. Depleting ATP levels reduces myosin stroke speed^48^, which leads to a lowering in the remodelling rate (larger *τ*_*rem*_) and increase in aster lifetime, finally jamming of the asters (*τ*_*rem*_ → ∞). In the ATP reduced state the remodelling rate of the aster has decreased by a factor of ∼4 (Supplementary Fig. S4e, i). Consistent with our predictions, the pattern changes from a mixed phase (Fig 4a-e) followed by a transient bull’s eye like pattern when the asters transition from a fast remodelling upon reduction of the aster remodelling state (Fig. 4g-I), in accordance with our predictions (Fig. 3b, c, see Supplementary Movie 5 and Supplementary Information-III Fig S2).

When ATP is completely depleted, with no detectable myosin activity Utr-wt remains clustered with a very slow decay in its intensity, Ezr-wt diffuses away (1000 s onwards (Fig. 4h-j, Supplementary Fig. S4j-p). This underlines the importance of continual myosin induced flows in driving the clustering of both the binders in the formation of the bull’s eye pattern. In the absence of myosin activity, diffusion restores the system back to its equilibrium distribution based solely on the binding affinities and without any spatial patterning within the aster. The high affinity of Utr-wt for F-actin explains why it remains clustered for an extended period.

#### Effect of relative density on steady state patterning

We next experimentally test the effect of relative density of the binders on active sorting (shown as blue box in Fig. 3c). This is achieved by having Utr-wt and Ezr-wt at different density ratios (n_Utr-wt_/n_Ezr-wt_) while maintaining the total linker concentration (Utr-wt + Ezr-wt) at the same level as in previous conditions (i. e., 20 nM). At n_Utr-wt_/n_Ezr-wt_ = 1/4, both Utr-wt and Ezr-wt exhibit active co-clustering, forming a mixed pattern in both the remodelling (Fig. 5a-d) and jammed aster states (Fig. 5e-h). This behaviour contrasts with the n_Utr-wt_/n_Ezr-wt_ = 1 scenario, where Ezr-wt is initially depleted from the aster core, forming a transient bull’s eye pattern, followed by a rapid reduction in local concentration when the aster jams (compare Fig. 5g with Fig. 4i; and Supplementary Fig. S5i with Supplementary Fig. S4k).

**Figure 5.**
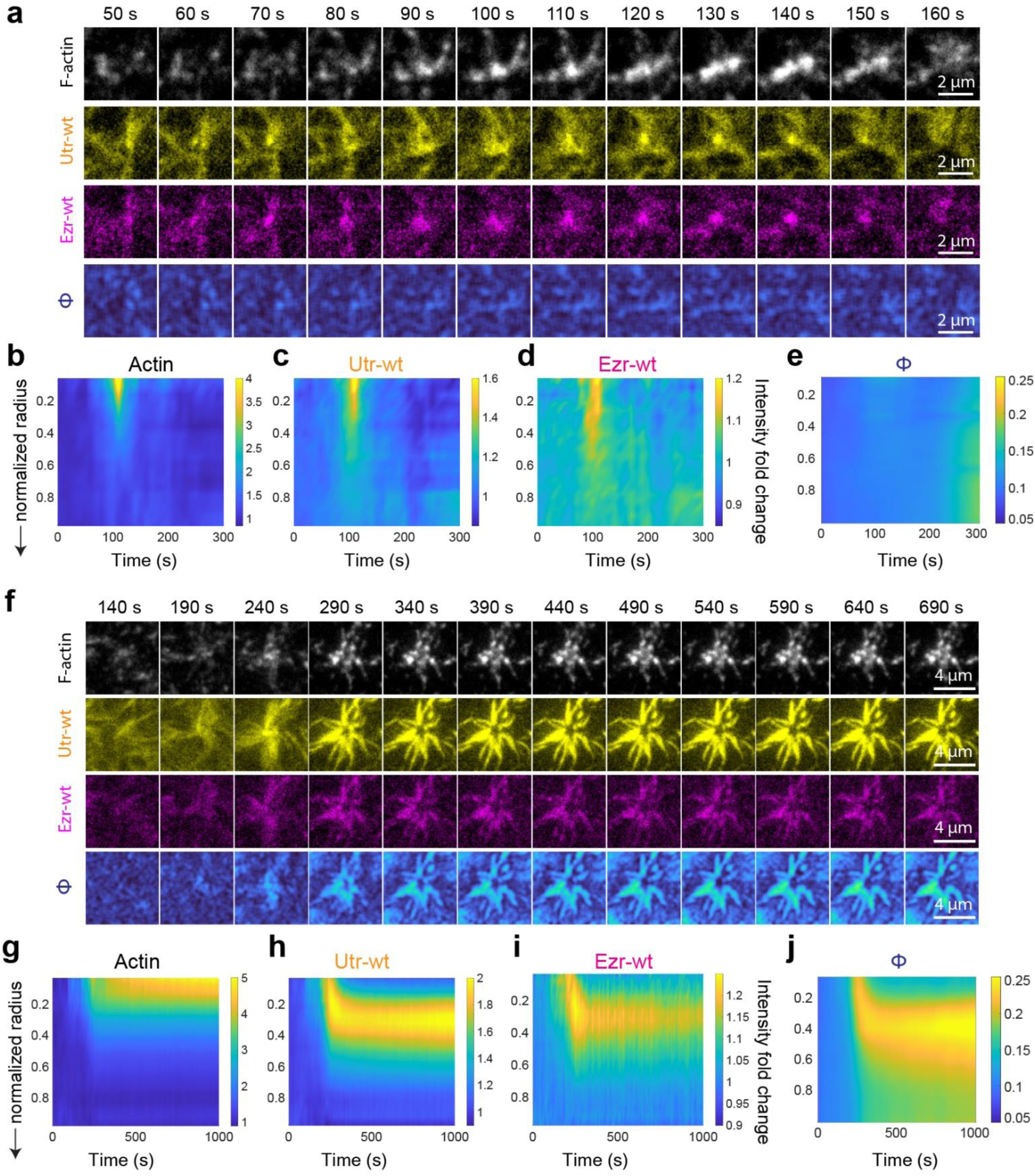
Effect of number density ratios on the sorting of binders. (**a-e**) Spatial organization of F-actin (grey), Utr-wt (yellow) and Ezr-wt (magenta) at ∼1/4 density ratio of Utr-wt/Ezr-wt in the remodelling state. (**a**) TIRF images show transient localized co-clustering of Utr-wt and Ezr-wt driven by contractile actomyosin flows (t ∼ 100 s; myosin-II added at t = 0 s), followed by dissipation of the clusters (t ∼ 130 s onwards). Surface plots show time-dependent radial intensity profiles of F-actin (b), Utr-wt (c), and Ezr-wt (d) and φ (e) (See Fig. S5 for more details) (n = 10 asters). (**f-j**) Spatial organization of F-actin, Utr-wt and Ezr-wt at ∼1/4 relative density during the jammed state. Both binders show robust prolonged co-clustering (t ∼ 390-1000 s), followed by dissipation of Ezr-wt (t ∼ 1000 s onwards), as shown by TIRF images (f) and Surface plots (g-j) (n = 19 asters). Scale bar, 2 μm in (a) and 4 μm in (f).

### Effect of protein-protein interactions on the stability of non-equilibrium patterns

There has been a lot of interest in proteins that exhibit monovalent and multivalent interactions amongst themselves, many of these are also known to bind to cortical actin directly or indirectly. Some prominent examples include CD44^1^, E-Cadherin^4^, T-Cell receptor, following the activation of LAT and recruitment of Nck^37^. Therefore, we explored the influence of on the strength of the attractive potentials between the proteins on this micro-phase segregation and the lifetime of the patterns in our simulations. We first start with having *K*_*A*_ > *K*_*B*_ and ∈_*AA*_= ∈_*BB*_, where the bull’s eye pattern with A in the inner region is obtained. We now tune ∈_*BB*_> ∈_*AA*_, keeping everything else fixed. We find that the bull’s eye parameter φk is enhanced. We also find that the strong attractive interactions have a strong influence on the lifetime of the patterns. This is demonstrated in **Fig. 3g** where we compare the time course of integrated densities of component A between two situations – when the attractive ∈_*AA*_ interactions are very weak (red curve, ∈_*AA*_= 0) and when they are much stronger (magenta, ∈_*AA*_= 1.5). We observe that the patterns formed in presence of the contractile platform dissolve quickly when the interactions are weak while those remain stable for longer periods even after the active dynamics are turned off.

### Active flows in cell membranes sort transmembrane proteins with varying actin binding affinities

To address the consequences of this active sorting mechanism in living cell membranes, we analysed membrane protrusions and retractions during lamellipodia formation in mouse embryonic fibroblasts (MEFs) spreading on fibronectin-coated glass^53^. In protruding cell regions, large scale retrograde (centripetal) flows of cortical actin emerge due to net actin polymerization at the leading edge of membrane protrusion and non-muscle myosin II driven contraction towards the cell centre^54,55^. We hypothesize that transmembrane proteins with ABDs of different affinities for actin will exhibit affinity-based sorting similar to our theoretical predictions and *in vitro* observations along this axis. Specifically, weaker actin binders should enrich towards the leading cell edge, while the stronger actin binders are accumulated towards the cell centre, along the axis of the retrograde flow of actin.

To test this idea, we express three transmembrane proteins in MEFs in three pairwise combinations (Fig. 6a). Each protein construct features an extracellular tag (GFP or Folate receptor (FR), a transmembrane (TM) domain and an ABD (Utr-wt, Ezr-wt, or Ezr-R579A) on the cytosolic side. The membrane proteins are fluorescently labelled with either anti-GFP nanobody (GFP-nanobody-Alexa546) or anti-FR Fab (MoV19Fab-Star635) and the cells are allowed to spread on fibronectin coated glass. Using TIRF microscopy, we track the spatio-temporal distribution of the fluorescently tagged transmembrane proteins during lamellipodia formation and retraction (Fig. 6b, c, see Supplementary Movie 6).

**Figure 6.**
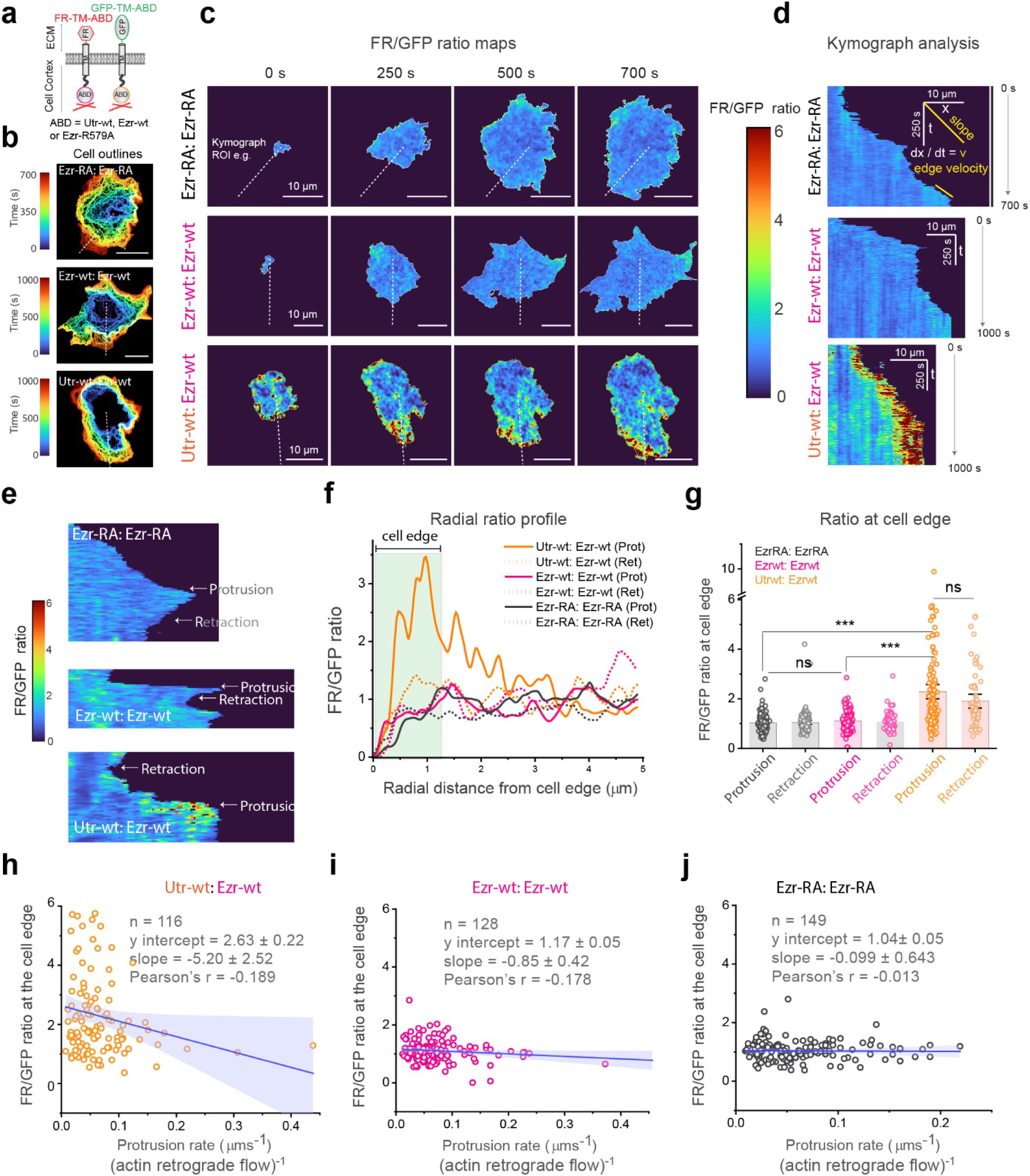
Retrograde actin flows at membrane protrusions are capable of sorting membrane proteins with differential actin affinities. **a,** Schematic of the constructs: two transmembrane proteins co- expressed in MEFs each with either a extracellular folate receptor domain labelled with a fluorescent folate analog (FR) or GFP fused to a cytosolic ABD from Utrophin (Utr-wt), wild type Ezrin (Ezr-wt) or non-binding mutant of Ezrin (Ezr-RA). **b**, Cell outline at successive time points (colour coded) show cell spreading dynamics for three binder combinations: Ezr-RA: Ezr-RA, Ezr-wt: Ezr-wt, and Utr-wt: Ezr-wt as indicated. **c**, Colour maps display FR/GFP intensity ratio at different time points for each binder combination. **d**, Kymographs generated from manually drawn line ROIs along membrane protrusions show both cell edge displacement and changes in FR/GFP ratio. Edge displacement speeds were used to calculate protrusion (or retraction) velocities, which were correlated to FR/GFP ratio at the cell edge (∼1.2 microns inwards). **e**, Representative kymograph close ups highlighting protrusion and retraction events and corresponding FR/GFP ratio for each binder combination. **f**, Line profiles of FR/GFP ratio along the protrusions and retractions (as in e), demonstrate sorting of FR- and GFP-tagged protein at the cell edge in the case of Utr-wt: Ezr-wt case. **g**, Distribution of FR/GFP ratio during protrusions and retractions for each condition (N= 3 independent experiments). Bars represent mean ± 95% confidence interval; (***) p<0.001; ns, non-significant (independent or paired t-test). **h-j**, scatter plots show the correlation between FR/GFP ratio and protrusion rate for Ezr-RA: Ezr-RA (h), Ezr-wt: Ezr-wt (i) and Utr-wt: Ezr-wt (j) pairs. A clear inverse correlation is observed only in the Utr-wt: Ezr-wt case. Scale bars, 10 μm (x) and 250 s (y).

We quantify sorting by calculating the spatial and temporal ratio of the two cell-surface proteins (see Methods) as exemplified in the kymograph analysis during lamellipodial dynamics (Fig. 6c-e, see also Fig. S6a-h). These kymographs allowed us to correlate instantaneous edge velocity with the profile of the ratio centripetally from the cell edge (Fig. 6e). Positive slopes correspond to protrusions and negative slopes to retractions where there is a consistent centripetal flow of actin towards the cell centre. The mean velocity distributions were comparable for the three conditions, ranging from ∼0.3 µm/s to −0.15 µm/s (Fig. S6i).

However, the Ezr-wt/Utr-wt ratio in each instance of extending edges decreased towards the cell centre, indicating sorting along the centripetal direction of the actin flow (Fig. 6f, g); Ezr-wt (the weaker binder) concentrates at the leading edge and Utr-wt (the stronger binder) is enriched towards the cell centre (Fig. 6f, g). This analysis indicates that the radial patterning is strongest at the edge, where the actin flows are fastest^56^, and requires differential F-actin binding, as exemplified by Utr-wt and Ezr-wt.

Furthermore, sorting corelates with protrusive edge velocity. In Utr-wt: Ezr-wt condition, the ratio at the edge shows a steep slope during protrusive velocities (Fig. 6i). This could be because retrograde actin flow correlates inversely with cell protrusion rate^57,58^. This corelation weakens when binder affinities are similar (Ezr-wt: Ezr-wt; Fig 6j) and is nearly absent when binding to the actin cytoskeleton is abolished (Ezr-RA: Ezr-RA^39 30^; Fig 6k). Sorting is diminished during retractive events (Fig S7j-i). The results are qualitatively consistent with our theoretical model, suggesting that cell membrane receptors coupling to active cortical actin flows result in their spatial and temporal sorting in living cells based on their relative affinity to F-actin.

## Discussion

The cell has many ways to generate protein clusters with specific compositions, for example by ordered assemblies in which proteins form homo- and heteromeric units. These assemblies reveal new interaction surfaces for creating multiple types of clusters^24^. We show here that in addition to equilibrium processes, cluster specificity/complexity may be achieved by differential binding affinities to the many components of the actomyosin cortex and their actomyosin remodelling rates.

Our experiments directly demonstrate how membrane proteins with different affinities for actin filaments result in precise spatial and temporal patterns. Central to this patterning are ATP-fuelled actin-flows and the relative affinity of the binders for dynamic actin. Our simulations predict three types of membrane protein patterns formed by contractile asters: *segregated*, *bull’s eye* and *mixed* as one varies the relative affinity and density of competing binders. In the mixed case, both the binders populate the contractile zone, while in the segregated case, the weaker binder is excluded from the contractile zone. In the bull’s eye pattern, the weaker binder forms an annulus around a cluster enriched in the stronger binder—the annulus becomes thinner as the pattern approaches the *segregated* phase. These features are reliably recapitulated in the experiments (Figs. 2, 4 and 5). The simulations also predict that increasing the relative actin affinity and density of the two binders improves sorting and patterning. Experiments using physiological actin-binding proteins, Utr-wt vs Ezr-wt (K_A_/K_B_ = 5) are in line with the predictions of our model.

We find that the sorting potential of asters enhances with their increased lifetime (Fig. 3f, Fig. 4f-j). This enhancement could arise from an active resonance phenomenon, as proposed earlier^5^, wherein the time scale of active advection of binders along dynamic F-actin aligns with the time scale of active aster remodelling (τ_rem_). The advection velocity of binders depends on their duty ratio for actin filaments, implying that longer aster lifetimes correspond to higher probability of differential advection leading to sorting.

In this study, the choice of the CH1-CH2 domain of Utrophin and C’ ERM ABD of Ezrin was made to represent the diversity of cellular actin-binding domains. The two domains were selected due to their distinct structural characteristics and F-actin binding mechanisms, with minimal spatial overlap on F-actin subunits. Additionally, these ABDs are ubiquitously used in membrane-cortex interactions by several membrane-cortex linkers^33,59–62^. It is also true that many cellular F-actin binders have spatially overlapping binding sites on F-actin that can lead to steric occlusion, especially if F-actin binding sites are limited^28,63,64^. This aspect of diversity is captured by Ezr-wt and Ezr-KA, which differ only by a single amino acid. However, despite potential overlaps, the overall spatial patterning is primarily driven by active advection and the relative affinities of the competing binders for advection.

Further we note that the contractile domains can potentially influence the local composition of functional molecules on the membrane is by driving the chemical purity of the clusters. Chemical purification could be achieved by in two different routes. First, through big differences in affinities to the domain of the two species i.e. *K*_*A*_/*K*_*B*_ is at one of the two extremes (≫ 1 *or* ≪ 1). If one of these conditions is satisfied, purity of the high-affinity species can increase with time. This is efficient even when the platforms do not remodel. However, if the two species are closely related, such as homologous proteins, with very similar affinities for the domain, the active remodelling feature of the actomyosin machinery can enhance the chemical purity of the cluster. Such purification process can be tracked by computing the local entropy of mixing in a given region of space, (defined as *S*_Ω_ = − ∑_*∝*_ *x*_*∝*_ ln *x*_*∝*_ where, *x*_*∝*_ is the mole fraction of component *α* in the given region Ω.). Subsequently, we can define the average chemical purity as Δ = 1 − 〈*S*_Ω_〉/ln 2, where the angular brackets represent averages over time and all chosen locations for the measurement. If the typical size of Ω is much smaller than the size of a contractile domain, such as 50-100 nm, such remodelling-driven purification is expected to be most effective (for example see Supplementary information-III, Fig. S3). Only a small number of molecules, like 5-10, can assemble in such a small area – which is close to the size of transient, actively-maintained nanoclusters observed in the cellular context. Therefore, this could provide the cell the ability to maintain chemical purity of these transient clusters for important biochemical reactions.

A plethora of plasma membrane-associated proteins and lipids can directly or indirectly associate with cortical actin, and our proposed mechanism provides a way for the living cell to spatiotemporally regulate its local composition through differential coupling to active flows in the cortex. Active contractile stresses can drive clustering of specific actin-binding components and can lead to the formation of distinct mesoscopic domains of specific and functional compositions in the membrane^9^. e.g., Outer leaflet GPI-anchored proteins form nanoscopic liquid-ordered domains due to their ability to participate in trans-bilayer coupling with active cortical stresses. These nanoclusters in turn build mesoscopic ‘active liquid-ordered emulsions’ recruiting via lateral interactions specific lipids in the plane of the membrane^1,9^.

Active clustering and micro-patterning also influence membrane-bound reactions in three ways: (i) reduce stochasticity through active local enrichment^25^, (ii) increase effective spatial coverage via discrete clustering^26^ and (iii) enhance signalling by sorting molecules in nano- and mesoscopic domains in space and time, excluding unnecessary cross-talk and inhibitors. e.g., during T-cell signalling, CD45 (a regulatory phosphatase) shows exclusion from the TCR-enriched core of the T-cell-APC immunological synapse^65–67^. While steric exclusion has been suggested as a dominant mechanism, driving micro-segregation of CD45 and TCR at the synaptic membranes, the role of differential F-actin binding and active sorting is worth exploring. It may be possible that formation of an active emulsion disfavours the access of CD45 phosphatase to the TCR-enriched signalling zone. Consistent with this a recent study using super-resolution showed that CD45 is pre-excluded form T-cell enriched actin rich microvilli in different lymphocytes^68^.

In summary, our study highlights the key role of active flows with sufficiently high Péclet numbers and membrane molecules with different duty ratios in driving active sorting phenomenon. Our *in vivo* results corroborate this idea, demonstrating that differential affinity for retrograde actin flows at the leading edge of spreading cells by *bonafide* actin-binding agents leads to spatial sorting of plasma membrane proteins. Specifically, the stronger binder aligns more along the flow towards the cell core, while the relatively weaker binder localizes predominantly at the leading edge. Further experiments are needed to elucidate the contribution of active flows at the cortex to the organisation and signalling of membrane proteins across various spatial and temporal scales.

## Methods

### In-vitro method

In this study, we used four different deca-Histidine tagged membrane-actin linker proteins by cloning and/or site-directed mutagenesis. First, we generated 10xHis-SNAP-EzrABD (Ezr-wt) by clonally appending the C’ actin binding domain of human Ezrin (EzrABD) to 10xHis-SNAP fragment^30,69^. Using site-directed mutagenesis, we next created two single amino-acid mutants of EzrABD: K577A and R577A2 to generate 10xHis-SNAP-EzrABD K577A (Ezr-KA) and 10xHis-SNAP-EzrABD R572A (Ezr-RA), respectively. Finally, we replaced the EzrABD fragment of 10xHis-SNAP-EzrABD with 261 amino acids from the N’ of Utrophin actin binding domain (UtrABD) to 10xHis-SNAP fragment to make 10xHis-SNAP-UtrABD (Utr-wt)^70^ (Fig. 1 and Fig. S2).

The linker proteins share a common set of features: a 10xHis (deca-histidine) tag for strong and specific binding to the Ni-NTA-DGS-doped bilayer membrane, a SNAP (or YFP) domain that allows specific fluorescent conjugation and subsequent visualization under the microscope, and a variable F-actin binding domain (ABD). The ABD is derived from the C-terminal ABD of human ezrin (Ezr-wt) (Fig. 1e), a point-mutant variant of the same (i.e. Ezr-K577A = Ezr-KA), or the N-terminal CH1-CH2 ABD of human utrophin (Utr-wt) (Fig. 1f).

Proteins were purified using Ni-affinity chromatography followed by size exclusion chromatography as described earlier. In some experiments, 10xHis-YFP-EzrABDwt was used instead of the SNAP-tagged version of Ezr-wt as indicated. Purified proteins were tagged with different SNAP-specific fluorescent dyes. Actin binding affinities were determined by (a) Actin co-sedimentation assay to determine their equilibrium affinities and (b) single molecule TIRF binding assay to determine their actin filament residence time ‘*t*’ (*t* =1/*k_off_*). How the proteins are prepared and how we assemble and image the system is all detailed in our recent publications^39,40,69,71^ (see Supplementary Information-II for details).

### Image analysis

Images saved in TIFF format were processed in Fiji (ImageJ, Open source) and Matlab (MathWorks, Academic License). Basic image analysis such as background subtraction, flat-field correction of TIRF images, multi-stack xy registration, photobleach correction, basic intensity measurement etc. is performed in Fiji. Further analysis was done in Matlab using custom-written scripts. Data are plotted in Matlab or Origin software. Different data statistics are estimated in Origin software (OriginLab Corporation, Licensed version) or Matlab. For more details, see *Supplementary Information* available online (see Supplementary Information-II for details).

### Segregation parameter analysis of in vitro data

To quantify microphase segregation of actin and actin-binding proteins *in vitro*, we computed a spatially resolved segregation parameter (φ) from time-lapse TIRF microscopy data. Image stacks for binder 1 and binder 2 were background-subtracted, flat-field corrected and corrected for any XY drift and photobleaching in Fiji and then imported as 16-bit TIFF files to Matlab and converted to double precision data. The intensity in each channel was normalized to pre-myosin intensity levels. A temporal rolling average (5-15 frame window) was applied to reduce noise and enhance detection of transient clustering. For each time point, local φ was calculated within a sliding 3×3-pixel spatial and a 5– 15 frame temporal window. The segregation parameter was defined as φ = I_A_ - I_B_/I_A_ + I_B_), where I is normalized intensity of A (stronger) or B (weaker) binder within the local window. φ maps were generated for each time point and exported as 16-bit TIFF stacks, using a fixed color scale for cross-frame comparison. In this context, φ = 0 indicates complete mixing, φ = 1 indicates complete phase separation, and intermediate values reflect partial segregation (see Supplementary Information-II for details).

### Theory and Kinetic Monte Carlo model

Here we present a brief description of the theory and the kinetic Monte-Carlo simulation that models our *in-vitro* experiments on the non-equilibrium clustering of two actin binding membrane proteins with different affinities towards F-actin. The current theory is primarily inspired by our previous active phase segregation picture in a multicomponent asymmetric bilayer membrane driven by the active contractile stresses^2,9,38,51^. Here we adopt the same theory using an off-lattice effective simulation model which sets it apart from the previous approaches.

In the off-lattice picture, we consider the supported bilayer membrane to be a 2D flat square surface with periodic boundary and side length *L*. On top of the surface, we put a binary mixture of two components A and B with number densities *n*_*A*_ and *n*_*B*_, respectively. Each protein molecule is represented by a coarse-grained sphere of diameter *σ*. The molecules interact among themselves via a modified Lennard-Jones potential where we fix the strength of repulsive interaction for any inter- or intra-species interactions. However, we tune the strength of homophilic attractive interactions which allow us to compare the features of clustering from equilibrium and with non-equilibrium origins. The existence of this equilibrium potential ensures that the molecules diffuse on the surface. We use the standard off-lattice Metropolis algorithm with a tunable displacement parameter to implement diffusive moves which obey detailed balance^72^. The number of diffusive moves per Monte-Carlo step, however, depends on the non-equilibrium moves described below.

Just below the above flat surface with the protein molecules, in a parallel juxtaposed layer we place the actomyosin fluid which applies localized contractile stresses on the molecules. As we considered earlier^9^, the stresses in these localized contractile regions are spatially correlated over a certain range and exhibit an exponential temporal correlation. Subsequently, we implement a Poisson birth-death process to realize the appearance and disappearance of these contractile regions. The remodeling times follow an exponential distribution with a characteristic timescale *τ*_*rem*_. When alive, any contractile domain radially advects any actin binding molecule within a range *ξ*, representing the range of spatial correlation, towards the core or the location of maximal stress. The binding-unbinding of the molecules to these domains is controlled by setting affinities expressed as duty ratios *K*_*A*_ and *K*_*B*_. In the simulation, the affinities and the advection speed are described by probabilities. In our model, *K*_*A*_ > *K*_*B*_ always and *n*_*A*_ ≤ *n*_*B*_. The advective moves are allowed based on a minimal displacement parameter. Such moves often incur a high energetic penalty and break detailed balance. We randomly allow such moves based on the advection probability if they are below the typical energy cost of hydrolyzing one molecule of ATP.

We also fix a Peclet number of 10 throughout our simulations to set the ratio of diffusive and advective moves. The detailed procedure of implementation of the simulation is provided in the SI appendix. Throughout our simulation, we fix the density of the contractile domains and ensure that only one such domain with size *ξ*∼ 10*σ* in present at any time.

We analyze the simulated configurations by calculating the time averaged local density difference *ϕ*(*r*) between A and B inside the contractile domain. For this we compute the temporally and spatially averaged radial densities *n*_*A*_(*r*) and *n*_*B*_(*r*) separately over a circular grid with spacing *σ*. Then we calculate the difference between these radial densities to find the average local density difference. From *ϕ*(*r*), we next compute the various Fourier amplitudes. The maximum amplitude, |*ϕ*(*q*)|_*max*_ is connected to the wave number *q* associated with the linear size of the largest domain observed in the system (Das PRL 2016). The larger the domain size of any one species within the region, the larger is the value of |*ϕ*(*q*)|_*max*_. We define arbitrary cut-offs on |*ϕ*(*q*)|_*max*_ values to distinguish among different patterns shown in Figure 3 (see Supplementary Information-III for details).

### Cell spreading assay

MEFs were transfected to express three different transmembrane proteins in pairwise combinations (see Fig. 7a) and fluorescently labelled at 37 °C for 15 min with either anti-GFP nanobody (GFP-nanobody-Alexa546) or anti-FR Fab (MoV19Fab-Star635). After washing with 1× M1 buffer (150 mM NaCl, 5mM KCl, 1mM CaCl2, 1mM MgCl2,20mM HEPES, pH 7.2) supplemented with 5mg/ml BSA cells were allowed to spread on fibronectin-coated glass (10ug/ml Human Plasma Fibronectin, Merck) and imaged using multicolour TIRF microscopy to monitor transmembrane protein organization during lamellipodia formation. Images were background-subtracted, and channels were aligned using custom MATLAB scripts based on cross-correlation of fiduciary markers. ROIs containing well-spread cells with high signal-to-noise were cropped and time-dependent drift was corrected using MultiStackReg Fiji plugin. Segmentation of cells was performed using the MovThresh module from the Edgeprops MATLAB pipeline, creating binary mask stacks to distinguish cells from background. Images were corrected for the minor cross-excitation of MoV19Fab-635 by the 561 nm laser, based on single-label controls; no significant GFPnb-546 bleed-through into the MoV19Fab-635 channel was detected. For each time point, normalized mean intensity values (from the segmented cell regions) were calculated and used to standardize pixel intensities, allowing comparison of relative protein enrichment or depletion, irrespective of absolute protein expression. Ratio images were consolidated by dividing the normalized images FR channel by those in the GFP channel (FR/GFP ratio), multiplied by a constant offset for visualization, and exported as 16-bit TIFF files. Using Fiji, ratio image stacks were used to manually draw ROIs along lamellipodia to generate kymographs. In Fiji, manually drawn line ROIs were used to track edge displacement over time, allowing us to calculate average edge velocities (positive for protrusions, negative for retractions). Edge intensity ratio was quantified within a 15-pixel (∼1.1 µm) radial distance from the cell periphery, enabling time-resolved analysis of protein localization and cell edge movement. To assess the difference in FR/GFP ratio at the cell edge during protrusion vs retraction events, we used paired or two sample t-test to evaluate statistical significance (p < .001). Scatter plots between FR/GFP ratio at the cell edge versus the edge velocity were plotted and analyzed by linear regression to calculate the y-intercept, slope and Pearson’s r.

## Supporting information

Supplementary Information-III

## Author contributions

Conceptualization: AB, AD, DK, MR, SM

Methodology: AB, AD, MI, SB, ST, DK, MR, SM

Investigation: AB, AD, MI, SB, ST, DK, MR, SM

Visualization: AB, AD

Funding acquisition: SM, MR

Project administration: DK, MR, SM

Supervision: DK, MR, SM

Writing – original draft: AB, AD, DK, MR, SM

Writing – review & editing: AB, AD, MI, SB, ST, DK, MR, SM

## Data and materials availability

All data are available in the main text or the supplementary materials.

## Acknowledgements

We thank all the members of the Mayor laboratory, in particular Kabir Hussain, Chaitra Prahakara, Parijat Sil and Bhagyashri Mahajan for suggestions and feedback on this manuscript. We thank the NCBS Central Imaging and Flow Facility as well as Amit Cherian and Greeshma Pradeep S for help with optical setups at NCBS. AB, SB and ST acknowledge doctoral fellowship support from National Centre for Biological Sciences, Tata Institute for fundamental Research (NCBS-TIFR).

## Supplementary Materials

Supplementary Figures S1-7

Supplementary Movies 1-6

Supplementary Information-I (Bioinformatics)

Supplementary Information-II (Experiments)

Supplementary Information-III (Theory and Simulation)

