## Supplementary Information-I for "Active flows drive clustering and sorting of membrane components with differential affinity to dynamic actin cytoskeleton"

### (Bioinformatics)

#### Identifying Actin-Associated Cell Membrane proteins

To get an estimate of the number and diversity of actin-binding proteins, we searched the UniProt database using the keyword "actin binding" (14th June 2022). From the results, we chose only the reviewed proteins from *Homo sapiens* and retrieved the Gene Ontology and subcellular localization details for this protein list (Figure 1) (Supplementary Data 1). We filtered the data further by checking for functional terms associated with actin, editing the misannotated entries and by manually adding missed out entries.

391 proteins were listed as experimentally verified actin-binding proteins, amounting to ~2% of all proteins in the cell (see **Table 1**). For the predicted actin-associated actin binding proteins in the cell, over 26% can associate with actin. Out of the 391 verified proteins, 18 were classified as integral membrane proteins associated with the cell membrane and 181 proteins were classified as peripheral (adaptor) cell membrane proteins, constituting 51% of total ABPs (**Fig. 1**).

To assign protein domain family (Pfam) to the listed ABPs, we scanned the protein sequences against the Pfam A library using a method already described in (Iyer MS et al, 2018). We identified 283 unique Pfam domains in these protein sequences, including 49 number of actin-binding domains. We annotated the actin-binding domains (ABDs) in the membrane-associated ABPs by referring to UniProt and literature mining (see **Table 2**). The most frequent ABD families in the membrane-associated ABPs has been indicated in the Fig. 1c. These include ERM_C and CH domains, the ABDs chosen for this study.

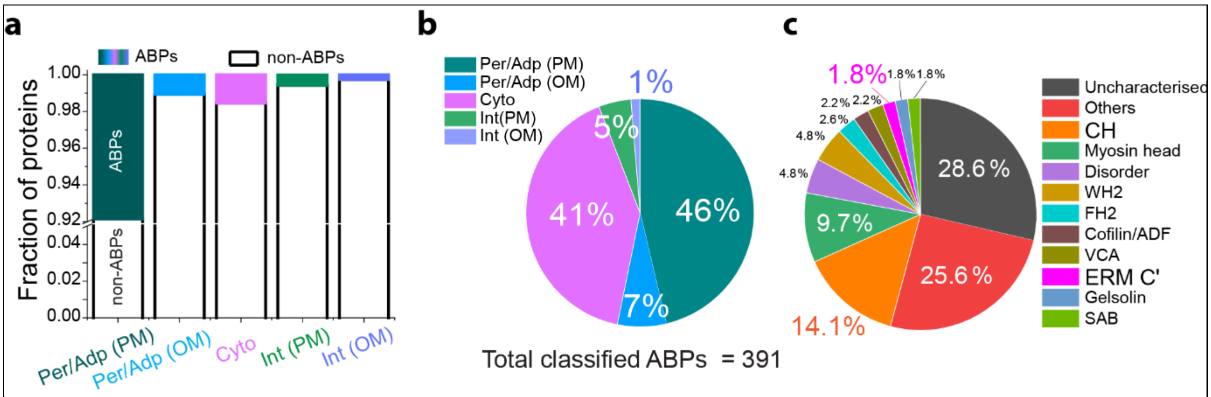


**Figure SI 1**: **The subcellular distribution of actin-binding proteins in the cell**. **a,** We classified actin-binding proteins with cell membrane association into integral (Int) or peripheral membrane proteins (Per/Adp). OM = other organellar membranes; Cyto = cytosolic. These proteins with actin-binding domains can directly link the plasms membrane (PM) to cortical F-actin. However, they can also undergo secondary interactions with cytosolic actin binding proteins (not included in this analysis). **b**, 8% of all peripheral proteins at the plasma-membrane [Per/Adp (PM)] and 0.5% of transmembrane proteins [Int(PM)] have experimentally verifiable ABDs, constituting up to 51% of cellular actin binding proteins listed in the analysis. **c**, Mapping of the actin-binding Pfam domains in the listed membrane-associated ABPs, the domains associating in high frequency have been plotted.

***Table 1*** **Categorization of cellular proteins based on actin-binding property and cellular localization.**

| **Category** | **Total proteins** | **ABP proteins (experimentally verified)** | **% of total fraction** | **AAP (predicted)** | **% of total fraction** |
|---|---|---|---|---|---|
| Cytoplasmic | 10348 | 160 | 1.5 | 1101 | 10.64 |
| Peripheral (Cell membrane) | 2305 | 181 | 7.8 | 224 | 4.34 |
| Peripheral (Organelle membrane) | 2550 | 27 | 1.05 | | |
| Integral (Cell membrane) | 3211 | 18 | 0.56 | 221 | 6.89 |
| Integral (Organelle membrane) | 1977 | 5 | 0.25 | 87 | 4.4 |
| **Total** | **20391** | **391** | **1.92** | **1633** | **26.27** |

*Table 2: Diversity of actin-binding domains in membrane-associated actin-binding proteins*

| Pfam domain | #Proteins with the domain |
|---|---|
| Uncharacterised | 65 |
| Others | 58 |
| CH | 32 |
| Myosin head | 22 |
| Disorder | 11 |
| WH2 | 11 |
| FH2 | 6 |
| Cofilin ADF | 5 |
| VCA | 5 |
| ERM_C | 4 |
| Gelsolin | 4 |
| SAB | 4 |
