## Supplementary Information-II for "Active flows drive clustering and sorting of membrane components with differential affinity to dynamic actin cytoskeleton"

**(Experiments)**

##### Cloning of 10xHis-SNAP-tagged Actin binding proteins

The four deca-Histidine-SNAP-tagged membrane-actin linker proteins used in this study were generated by site-directed mutagenesis and cloning in pET28a vector backbone. First, 10xHis-SNAP-EzrABD (Ezr-wt) was cloned by appending the C' terminal actin binding domain of human Ezrin (EzrABD) to 10xHis-SNAP fragment inside the pET28a vector (Saleh et al., 2009; Shrivastava et al., 2015). Next, by site-directed mutagenesis in EzrABD of Ezr-wt construct, two single-amino-acid mutant versions of Ezr-wt were created: 10xHis-SNAP-EzrABD K577A (Ezr-KA) and 10xHis-SNAP-EzrABD R579A (Ezr-RA), respectively. Finally, the EzrABD fragment of Ezr-wt was replaced with 261 amino-acid long N' terminal actin binding domain of human Utrophin (UtrABD) (Burkel et al., 2007) to make 10xHis-SNAP-UtrABD (Utr-wt). In some experiments we used 10xHis-YFP-EzrABDwt (Shrivastava et al., 2015) instead of the SNAP-tagged version of the Ezr-wt and that has been clearly mentioned in the corresponding figure legends. All four constructs were confirmed by Sanger sequencing.

##### Expression and Purification of the SNAP-tagged Actin binding proteins

BL21(DE3)* E. coli were transformed with 50-100 ng of pET28a-10xHis-SNAP-ABD plasmid DNA and plated overnight on LB agar containing 50 µg/ml kanamycin at 37 °C. A single colony was picked and inoculated in 50–100 ml primary LB culture (50 µg/ml Kanamycin), which was grown at 37 °C until the OD600 reached 0.5–0.8 (typically takes 5–6 h). Approximately 20–25 ml of this culture was used to seed each 500 ml secondary culture in 2 L baffled flasks containing LB (50 µg/ml Kanamycin), which were grown at 37 °C with shaking at 200 rpm until the OD600 reached a value of 0.7–0.9 (typically 3–4 h). Cultures were cooled to ~20 °C for 10–15 min before induction with 200 µM IPTG, and cells were grown overnight at 18 °C (alternatively, cultures were induced at room temperature and grown at 37 °C for 3–4 h). Cells were harvested by centrifugation at 5000 × g for 10 min at 4 °C, and pellets were either processed immediately for lysis or flash-frozen in 1x PBS supplemented with 0.1–1 mM PMSF for storage at –80 °C.

Cell pellets obtained from 1 L bacterial culture were resuspended in 40–50 ml of lysis buffer and sonicated in a 100 mL glass beaker using a probe Sonicator at 35% amplitude, 10 s ON, 10 s OFF, pulsed for 15–30 cycles, totaling 5-10 min. Lysate was clarified by centrifugation at 20,000 × g for 30 min at 4 °C and passed through a sterile 0.45 µm filter before loading onto a 5 ml HisTrap FF Excel column (Cytiva) pre-equilibrated with binding buffer, at 0.5 ml/min, using a peristaltic pump. After washing with 10 column volumes (CV) of wash buffer at 1 ml/min, the column was connected to an ÄKTA system for elution and automatic fractionation. Elution was carried out using a four-step protocol: (i) an additional 5 CV wash, (ii) a linear imidazole gradient from 20–40 mM to 500 mM in 10 CV, (iii) 5 CV of 100% high-imidazole buffer, and (iv) re-equilibration with 5–10 CV of wash buffer. Eluted fractions (0.5–1 ml) were collected and analyzed by SDS–PAGE, and those containing the target protein were pooled. Protein concentrations were estimated by comparison to BSA standards on SDS–PAGE, by A280 nm absorbance, and by Bradford assay.

When necessary, pooled protein fractions were spin-concentrated using centrifugal concentrators (3–10 kDa MWCO, Millipore or Pall), ensuring concentrations did not exceed 5 mg/ml to avoid aggregation. The concentrated protein was further purified by size-exclusion chromatography (SEC) on a Superdex 75 or Superdex 200 column (GE Healthcare) pre-equilibrated with gel-filtration buffer using an ÄKTA system. Proteins were injected in volumes of 150–500 µl (Superdex 75 10/300) or 0.5–4 ml (HiLoad Superdex 75/200 16/600), and fractions were collected during the main peak elution. SDS–PAGE was used to confirm protein purity, and pure fractions were pooled and quantified for concentration. For long-term storage, purified proteins were supplemented with 10–20% sterile glycerol, aliquoted, flash-frozen in liquid nitrogen, and stored at –80 °C for longer shelf life.

**SNAP-labelling of purified 10xHis-SNAP Actin Binding proteins:**

Frozen aliquots of SNAP-tag fusion proteins were quick-thawed at 37 °C, clarified by ultra-centrifugation at 250,000 × g for 15 min at 4 °C, and the supernatant was supplemented with fresh TCEP to a final concentration of 2–5 mM. Protein samples were mixed with SNAP-reactive dyes (e.g. Benzyl Guanine-based fluorophores) at a 2–3 molar excess, ensuring DMSO concentrations remained below 1% to avoid reduced labeling efficiency. Typical reactions contained 135.8 µl of 15 µM protein and 4.2 µl of 1 mM dye (~2-fold molar excess) in a final volume of 140 µl, which is compatible with desalting using a PD SpinTrap G-25 column. Labeling reactions were incubated for 30 min at 37 °C, or 1 h at room temperature, in the dark.

Following incubation, free dye and protein aggregates were removed either by size-exclusion chromatography (SEC) on a Superdex-75 10/300 column, or by PD SpinTrap G-25 desalting columns (Cytiva) equilibrated with labeling buffer (50 mM Tris-HCl pH 7.5, 150 mM NaCl, 5 mM TCEP, 0.1% Tween-20). For SpinTrap purification, columns were equilibrated by five washes with 400 µl buffer (spun at 800 × g for 1 min each time) to remove stabilizers and preequilibrated the column before loading 100–180 µl of sample. Labeled protein was recovered by centrifugation at 800 × g for 2 min. For higher-volume reactions (up to 2.5 ml), PD-10 desalting columns were used following the manufacturer's spin or gravity-flow protocols. In all cases, eluates were collected, and protein concentration and labeling efficiency were determined spectrophotometrically by measuring absorbance at A280 for protein and at the dye-specific absorbance maximum, using correction factors (CF280 nm) provided by the dye manufacturer or determined empirically using the pure dye solution.

Labeling reactions were generally robust across a range of pH (7.0–8.0), salt (50–350 mM NaCl), and detergent conditions (0.05–0.1% Tween-20, which does not impair SNAP activity). SNAP-tag stability and labeling efficiency were enhanced by reducing agents (TCEP or DTT), while chelating agents such as EDTA were avoided to preserve the $Zn^{2+}$ cofactor for SNAP proteins. For long-term storage, labeled proteins were supplemented with 10–20% glycerol, aliquoted in 2–10 µl volumes, flash frozen in liquid nitrogen, and stored at –80 °C for up to two years.

**Buffer Compositions for HSE Protein Purification**

Lysis Buffer (pH 7.5): 50 mM Tris-HCl, 150 mM NaCl, 1–5 mM TCEP (note: TCEP is unstable in phosphate buffers), 0.1% Tween-20, 0 mM EDTA (excluded to avoid chelation of the SNAP-tag's structural $Zn^{2+}$), 20 mM Imidazole, and 0.1–1 mM PMSF and/or 1 cOmplete™ protease inhibitor tablet per 50 ml.

Binding Buffer (pH 7.5), for $Ni^{2+}$ Affinity Chromatography: 50 mM Tris-HCl, 500 mM NaCl, 1–5 mM TCEP, 0.1% Tween-20, 20 mM Imidazole.

Wash Buffer (pH 7.25): Same as Binding Buffer, but with 40 mM Imidazole. For HSEzr-R579A variant, Binding Buffer (20 mM Imidazole) can be used as Wash Buffer.

Elution Buffer (pH 7.0): Same as Binding Buffer, but with 500 mM Imidazole.

Gel Filtration Buffer (pH 7.5): 50 mM Tris-HCl, 150 mM NaCl, 5 mM TCEP (again, avoid phosphate buffer with TCEP), 0.1% Tween-20.

All buffers were filtered through a 0.22 µm membrane and degassed by vacuum, sonication, or centrifugation at 10,000 × g for 20 min prior to use.

**Purification and labelling of Actin, myosin and capping proteins:**

Skeletal actin, myosin -II were extracted from fresh chicken skeletal muscle and purified as described in detail in our previous publication, (Köster et al., 2022, 2016).

**Purification and labelling of capping protein**

Murine capping protein-- a heterodimer of α1 and β2 subunits-- was purified following an earlier protocol (Bieling et al., 2018; Funk et al., 2021). First, pETM20 (containing Trx-6xHis-TEV-α1-subunit) and pETM33 (containing 6xHis-GS-PreScission-SNAP-β2-subunit) were co-expressed in Rosetta DE3 E. coli. Cells were grown at 37 °C in shaking LB media with Ampicillin, Kanamycin and Chloramphenicol antibiotics till OD595 = 0.6-0.9. The protein expression was then induced with 250 µM IPTG and cells were grown at 18 °C for 10-12 hrs. Next, the cells were pelleted at 5,000 x g for 10 min and lysed by sonication in the presence of Lysis buffer on ice (Sonics VC750, tip diameter = 12 mm) using the same setting as described for HSE purification. The lysate was spun at 20, 000 x g for 20 minutes to remove cell debris (Beckman Coulter Avanti J-26 XP). The supernatant containing the protein was passed over a $Ni^{2+}$ affinity column and eluted with 400 mM imidazole. That was followed by combined TEV/PreScission cleavage on ice for 12-14 hrs. The cleaved protein mix was then passed over a GE HiLoad Desalting column to remove imidazole. The uncleaved protein, free tags, and the His-tagged Protease fragments were removed by re-circulating the protein mix through the $Ni^{2+}$ affinity column. The fully cleaved heterodimer from the flow-through was further purified by passing it over an anion-exchange Mono Q column (GE Healthcare). The purified protein was labeled with a 1.5-2-fold molar excess of SNAP-Surface AF-647 (NEB), overnight on ice. The labeled protein was separated from unreacted dye by gel filtration on a GE Superdex-200 column. Protein purity level was determined by SDS-PAGE; concentration and degree of labelling were measure by spectrophotometry using $A_{280nm}$ of capping protein ($\varepsilon_{280nm}$ = 99,530 $M^{-1}cm^{-1}$) and $A_{\lambda max}$ of the dye. Purified capping protein was aliquoted, snap-frozen in liquid N2 in Capping protein storage buffer and stored at -80 °C. The activity of the labelled capping protein was checked in-vitro: fluorescent fixed amounts of G-actin (10% labelled) were polymerized in the presence of different capping protein amounts. The filaments were imaged under TIRF microscopy, and the length distribution of fluorescent actin filaments was quantified. The higher the relative concentration of capping protein, the shorter was the filament length distribution (Köster et al., 2022, 2016).

**Actin affinity measurements**

Three different assays were performed to measure the actin-binding affinity of the membrane-actin linker proteins: F-actin co-sedimentation assay, Single-molecule dwell time assay, and FRAP in the presence of SLB bound F-actin. The FRAP assay will be discussed in section 2.10.

**F-actin cosedimentation assay**

This classical biochemical assay yields the equilibrium dissociation constant (K_D) of actin-binding proteins. In this assay, different amounts of the actin-binding protein (ABP) of interest are incubated with a fixed amount of F-actin. The reaction mixes are then spun at ultrafast speed to separate the actin-bound and free fractions of ABP. The extent to which the ABP co-sediments with F-actin is directly proportional to its equilibrium actin-binding affinity (K_D) of the protein.

First, G-actin and ABP were spun at 110,000 x g for 20 min at 4 ºC to remove any protein aggregates. G-actin was then polymerized at 5-10 µM final concentration and fixed amounts of actin filaments were added to TLA 100.3 ultracentrifuge tubes (with maximum volume = 150 µL). Total number of ultracentrifuge tubes required depends on the number of data points one intends to cover plus two control tubes. The final concentration of actin was kept same in all the tubes; except no actin was added to the ABP-Only control tube (C2).

Final actin concentration typically ranges from 0.1-5 µM (in terms of G-actin) and depends on the expected affinity of the ABP being tested. For F-actin concentrations lower than 0.5 µM — the critical concentration of actin—actin filaments were stabilized with phalloidin at 2:1 molar ratio (actin: phalloidin). Ideally, the final F-actin concentration selected should be 5 to 10 times lower than the expected $K_D$ of the ABP (Pollard, 2010).

Next, the concentration of the pre-spun ABP was determined with A280nm and/or Bradford assay. Different amounts of ABP were added to each tube, typically ranging from 0.01-15 µM, except no ABP was added in the Actin-Only control tube (C1). Concentration range and step size is determined by the amount of F-actin added in the tubes. The final reaction volume of each tube was made up to 75 µL with 1x KMEH buffer.

The samples were mixed with gentle pipetting and 1/3 of the volume was taken out from each tube as "total" fraction (to generate a calibration curve later). The remaining 50 µL were equilibrated for 30 min at RT and then spun at 110,000 x g for 20 min at 25 ºC. Carefully, the "supernatant" from each tube was removed and kept. The "pellet" in each tube was resuspended in 50 µL of G-buffer. The fractions were resolved by SDS-PAGE using 10% polyacrylamide gels, stained with 0.01% Coomassie brilliant blue, and digitized with an EMCCD camera on ImageQuant (GE Healthcare).

**Single-molecule dwell-time assay**

This single-molecule biophysical assay allows us to measure the dwell time (${k_{off}}^{-1}$) of single ABP molecules once they have bound to an actin filament. The dwell time is inversely proportional to the dissociation rate constant, $k_{off}$, of the ABP.

In this assay, a flow-cell was prepared by sticking an ultraclean glass coverslip on a clean microscope slide with a double-sided tape. Usually, three lanes were made per coverslip, and one lane was used at a time. Solutions were flowed in from one side and a tissue was used to create a capillary force on the other side to give a flow to the fluid. First, the coverslip surface in each lane was coated with 10 µL of BSA-Biotin (1 mg/ mL) followed by addition of 10 µL Streptavidin (1 mg/ mL). Unbound Streptavidin were washed out with the wash buffer and the surface was passivated with BSA (10 mg/ mL) for 5 min. Next, fluorescent actin filaments coated with Biotin-xx-Phalloidin were flowed in and allowed to specifically bind to the surface-bound Streptavidin. Next, the flow-cell was mounted on the microscope and intensity of F-actin was checked. Imaging was always done in the middle of the lane. Subsequently, fluorescent ABP was added at picomolar ($10^{-12}$ M) concentration.

TIRF imaging was done to capture the single binding/unbinding events. For each region of interest (typically 25x25 $\mu m^2$), three to five images of F-actin were recorded in the beginning. Next, the ABP was imaged @ 10-20 Hz for up to 100 s. To improve the lifetime of fluorophores, imaging was done in the presence of an oxygen-scavenging complex (PCA/PCD) and a triplet-state inhibitor (Trolox). Images were saved in 16-bit TIFF format.

**Sample Preparation for In-vitro Experiments**

**Cleaning glassware**

Freshly cleaned No. 1 Borosilicate coverslips (round or rectangular) were used to prepare samples for the in-vitro experiments. Clean amber glass vials (Thermo Scientific or Supelco) were used to store and mix chloroform solutions of lipids. Glassware was cleaned by bath sonication at 65°C for 20-30 min (Optics Technology). First, the glassware was treated with

Hellmanex II or III (Hellma Analytics) followed by a thorough rinse with MilliQ water. Next, the glassware was treated with 1-2N NaOH followed by another thorough rinse with MilliQ water. The coverslips were kept in MilliQ inside a Coplin jar for up to 6 hours and dried under $N_2$ gas stream immediately before the experiment. The glass vials were dried inside a hot air oven at 60°C and used within 6 weeks.

**Making sample chambers**

Sample chambers were prepared by sticking half-cut PCR tubes to freshly cleaned glass coverslips as described earlier (Köster et al., 2016). First, lids and the lower conical halves of autoclaved PCR tubes (Tarsons) were cut out with a sharp surgical blade. The cylindrical upper halves were stuck inverted to clean coverslips using a UV curable adhesive (NOA88-8801-Norland Products) such that the uncut side with smooth edges touched the coverslip. The rectangular coverslips accommodated upto three reaction chambers; the round ones accommodated only one. The chamber-containing coverslips were UV-illuminated for 3-5 min under vacuum. The coverslips were then taken out and each chamber was individually tested for leakage with MilliQ water. Each chamber could hold up to ~130 µl of sample. For imaging, the sample was mounted on the microscope stage using either a custom-designed coverslip holder (for rectangular coverslips) or an Attofluor™ cell chamber from Thermo Fischer (for circular coverslips).

**Preparing small unilamellar vesicles**

Multi-lamellar vesicles (MLVs) of fixed compositions were prepared by mixing various lipids in clean amber glass vials. Typically, chloroform solutions of DOPC and DGS-NTA-$Ni^{2+}$ lipids were mixed at 98:2 mol%. The mixture was dried under a slow stream of $N_2$ gas. The dried lipid film was vacuum-desiccated for >2 hrs to remove any traces of chloroform. The desiccated lipid mix was resuspended in SUV rehydration buffer (50mM HEPES, 150mM NaCl, 5% sucrose, pH 7.5) to make a 4 mM suspension of multi-lamellar vesicles (MLVs). The vesicles were aliquoted in 1.5 ml eppendorf tubes (Tarsons) and layered with Argon gas to minimize oxidation. The aliquots were flash-frozen in liquid $N_2$ and stored at -20°C for up to 6 weeks.

Small unilamellar vesicles (SUVs) were made by repeated freeze-thawing of MLVs: the MLVs contained in 1.5 ml Eppendorf tubes were flash-frozen in liquid $N_2$ for 10-20 sec immediately followed by thawing at 40-45°C in a water bath; the cycle was repeated 15-20 times until the solution looked less turbid. The solution was then passed through an extruder (Avanti Polar lipids) with an 80nm pore-size polycarbonate filter membrane (GE Whatman) to form monodispersed SUVs of ~100nm diameter. Alternatively, the solution was microtip-sonicated with the following settings: 10 secs ON, 60 secs OFF on ice, 30% of maximum amplitude, 3-5 cycles until the solution turned visibly clear. To remove any lipid debris, the sonicated suspension was centrifuged at 20,000 x g for 60 min at 4°C. 80% of the supernatant, containing the SUVs, was collected and stored on ice. The shelf-life of SUVs depends on their composition— DOPC: DGS-NTA-$Ni^{2+}$ SUVs were stable for up to 6 days.

**Making supported lipid bilayers**

Supported lipid bilayers (SLBs) were formed by adding SUVs to fresh, hydrophilic sample chambers. Each sample chamber was pre-hydrated with 100 µl of SLB formation buffer (50 mM Hepes, 150 mM NaCl, 2 mM CaCl2, pH 5.5 or 1x PBS, pH 5.5). 8-15 µl of DOPC: DGS-NTA-$Ni^{2+}$ SUVs were incubated for 15-20 min at RT. During this time, the dense SUVs containing 5% sucrose adsorb to the hydrophilic glass bottom. The vesicles rupture via vesicle-surface and vesicle-vesicle interactions, forming bilayer patches, which upon reaching a critical area coverage propagate like a wave and coalesce into a continuous planar bilayer (Andrecka et al., 2013). The unfused lipids were removed by 10 -12 gentle washes with SLB storage buffer (50 mM Hepes, 150 mM NaCl, pH 7.2). β-Casein (0.1mg/ml) was added to block bare glass patches.

**Assessing quality of lipid bilayers**

The quality of supported lipid bilayers was assessed by fluorescence recovery after photobleaching (FRAP). A fluorescent Poly-His-tagged protein (10-15 nM) was incubated on β-casein passivated lipid bilayers for 30-45 min. Unbound protein was removed by multiple washes with 1x KMEH buffer (50 mM KCl, 2 mM MgCl2, 1 mM EGTA, 20 mM Hepes, pH 7.2). The fluorescence intensity of the bilayer-bound fluorophore was checked with total internal reflection fluorescence (TIRF) microscopy. Good quality bilayers show a large-scale, uniform distribution of the fluorescence intensity. Bad bilayers show intense and patchy fluorescence spots. The integrity of the bilayers was further confirmed by FRAP. A high intensity laser beam was concentrated on a small circular region of the bilayer—by closing the field diaphragm of the microscope—to locally bleach all the fluorophores. The field diaphragm was then reopened and the recovery of fluorescent signal in the bleached spot was recorded under TIRF. Good bilayers showed a consistently fast diffusion coefficient: D = 0.8-1.5 $um^2s^{-1}$ for poly-His tagged fluorescent proteins (or D = 1.5-3 $um^2s^{-1}$ if using a fluorescent lipid, such as Rhodamine-PE, as a bilayer mobility probe). Bilayers with immobile or slowly diffusing fluorophores were discarded.

**Hierarchical assembly of an active actin-membrane composite with defined protein densities.**

The process begins with the formation of a fluid SLB composed of DOPC and Ni-NTA-DGS (98: 2 mol%) on a clean, hydrophilic cover-glass, as described earlier. Next, the sample surface is passivated with BSA or β-Casein to block any bare regions of the glass substrate. The membrane-actin linker proteins are added at defined concentrations and allowed to equilibrate on the lipid membrane. The unbound protein is washed out and the sample is mounted on the TIRF microscope and time-lapse imaging is initiated.

To incorporate actin-binding proteins (ABPs) at controlled surface densities, fluorescent 10xHis-SNAP–tagged proteins were added to β-casein–passivated bilayers to reach a total binder concentration of 20 nM, unless otherwise mentioned. For example, in stoichiometric conditions (nA/nB = 1), 10 nM each of Ezr-wt and Ezr-KA were incubated on the bilayer in a total reaction volume of 100 uL for 30–45 min at room temperature, followed by 3x washes and then the addition of F-actin, myosin-II, and ATP (Köster et al., 2022). For other ratios, such as nA/nB = 0.25, 5 nM Utr-wt and 15 nM Ezr-wt were used prior to washing of free binders followed by addition of capped F-actin.

Next, a known amount of pre-polymerized, fluorescent actin filaments was added to the membrane. For polymerization, dark and fluorescently labelled G-actin are mixed at 10:1 ratio and then co-polymerised in the presence of murine CapZ protein to obtain filaments of controlled length. Carefully, F-actin was added with a blunt-ended pipette tip to the bilayer to avoid shearing and allowed to incubate for 20-30 min.

After adding F-actin, the bilayer was scanned under the microscope to check the spatial distribution of HYE and actin filaments. An artifact-free region of interest, with uniform intensity of HYE and F-actin, was selected for subsequent imaging of the sample. After adjusting the imaging conditions, such as the laser power, integration time etc., a time lapse recording was initiated. Typically, 10 to 30 images of the selected field of view were acquired (time interval between two successive frames is 5-15 s). These 'pre-myosin' images are used to normalize the intensity data in each channel. After that the time-lapse recording was paused. A known amount of myosin II (along with oxygen-scavenging PCA/PCD mix, ATP regeneration mix etc.) was carefully added to the system without disturbing the sample. The time-lapse recording was resumed and continued for up to 120 minutes.

The system has a limited amount of ATP which is consumed by the actomyosin activity. Eventually, the actomyosin asters are arrested in a ATP-depleted jammed state (described ahead). One can add more ATP (0.1-1 mM) to the system and initiate a new cycle of myosin contraction. Usually, the actomyosin dynamics during this cycle is faster than the first cycle (likely because myosin heads are already engaged with membrane-bound F-actin), hence the time-lapse recording was done at a faster frame rate (time interval is 2-5 s) and usually for 15-30 min. Images were saved in TIFF format along with metadata. Image analysis was done in ImageJ and Matlab (MathWorks).

**Image Processing and data analysis**

All images were saved or converted into *tiff* format and processed in Fiji (ImageJ) and/or Matlab. Basic image analysis such as background subtraction, flat-field correction of TIRF images, multi-stack registration, photobleaching correction, intensity measurement etc. was performed in Fiji. Further analysis was done in Matlab using custom-written scripts. Data were plotted in Matlab or Origin software. Different data statistics were estimated in Origin software.

**TIRF FRAP analysis**

The fluorescence recovery after photobleaching (FRAP) assay was used to assess the integrity of fluorescent supported lipid bilayers. Time-lapse movies were recorded encompassing pre-beach fluorescence intensity and the post-bleach fluorescence recovery. Images were processed in Fiji. After background subtraction, mean intensity values were measured from the bleached spot and a reference region. The recovery data were exported to Origin software. The recovery profile was fitted to an exponential model to determine the characteristic recovery time and $t_{1/2}$ from which the diffusion coefficient, D, was calculated.

**Co-sedimentation assay analysis**

To measure equilibrium F-actin affinity with this assay, different amounts of actin-binding protein were incubated with a fixed amount of F-actin. Actin-bound and unbound fractions of the proteins were separated by centrifugation followed by SDS-PAGE. Gels were scanned with ImageQuant (GE) and data were saved as 16-bit *.tiff* images. Using Fiji, ROIs were drawn on the images to measure integrated densities from protein bands and background regions. Data were imported to Matlab in .csv format. A calibration curve was generated to convert the band densities into known protein amounts. Actin-bound fractions of the actin-binding protein were calculated at different actin to protein ratios. An isotherm was plotted between the actin bound fractions of ABP (mol/mol of actin added) and the free ABP added ($[L]$). The curve was fitted to Hill's equation:

$$\frac{{B_{max}}^n + [L]^n}{[L]^n + {K_d}^n}$$

where, $K_d$ is the equilibrium dissociation constant, $B_{max}$ is the binding stoichiometry, and $n$ is the binding cooperativity.

**Single-molecule dwell time analysis**

Fluorescent actin was immobilized on clean, passivated coverslips covered with the assay buffer. Fluorescent actin binding proteins were added in pM concentration. Single binding and unbinding events were recorded with TIRF microscopy. The data were saved as 16-bit .tiff image sequences. Using Fiji, the images were converted into kymographs, with binding events showing as bright lines. Using a custom-written Matlab script, the Kymographs were thresholded to remove single-pixel-long noisy events. The length of the remaining lines was measured and converted back into time durations (1 pixel = 100msec). The data were plotted as semi-log frequency plots and fitted to an exponential model to determine the characteristic dwell time (${k_{off}}^{-1}$) of each actin binding protein.

**Microscope Instrumentation and Image Acquisition**

Nikon Ti Eclipse microscope was used for high-resolution, multicolour TIRF and Widefield microscopy. The microscope was equipped with a TIRF unit connected through a polarization-conserving optical fibre to an Agilent monolithic laser combiner MLC400 with four laser lines: 405, 488, 561, and 640 nm (Agilent Technologies), two motorized turrets, multiple objectives, a motorized stage, and a Perfect Focus System. Each laser line was coherent, monochromatic and plane polarized. The upper turret was fitted with a quadpass dichroic to direct the excitation beam from the optical fibre and TIRF illuminator towards the sample through the Objective. The lower turret was fitted with different emission filters for sequential, multicolour imaging. Images were collected with a 100X, 1.45 NA Nikon Oil Objective with a 512 x 512-pixel EMCCD camera (Photometrics Evolve 512) yielding a pixel size of 153 nm (or 103 nm when a 1.5x tube lens was inserted before the camera), or 1200 x 1200-pixel sCMOS cameras (Photometrics 95B primer) with a pixel size of 102 nm (73 nm with 1.5x tube lens).

In TIRF illumination mode, the laser light was focused on the back-focal plane of the high NA objective with a motorized mirror, away from the central axis of the objective, to create an oblique supercritical-angled total internal reflection of the excitation beam. This results in an evanescence field at the glass-water interface, which decays exponentially from the surface along the z-axis. The evanescent field is wavelength specific and can excite only the fluorophores at or close enough (~ 150 nm) to the glass surface (Axelrod, 2008). In Widefield mode, the beam was focused on the back-focal plane of the objective, right along the central axis, to create a uniform Köhler Widefield illumination through the sample. Single camera detection mode was used for simple TIRF and Widefield intensity measurements.

For TIRF anisotropy measurements, the polarization of the excitation laser was further purified with a clean-up polarizer to obtain an extinction coefficient (PA/PE) of ~1500:1. This plane polarized light was propagated through a quadpass dichroic to the high NA objective to hit the substrate surface at a supercritical angle such that it created an s-polarized evanescence field (i.e. the electrical (polarization) vector of the light is orthogonal to the plane of incidence). Unlike p-polarized evanescence, s-polarized evanescence creates a purely polarized field in the xy plane that decays exponentially from the surface in the z direction (Axelrod, 2008, 1989; Ghosh et al., 2012; Lakowicz, 2006). The emission from the sample was directed as a collimated beam to the two cameras, aligned and positioned orthogonal to each other, with the help of a Moxtek polarizing beam splitter. The splitter transmits the p-polarized component of the emitted light through to the *Pa* camera and simultaneously reflects the s-polarized component to the *Pe* camera. These *Pa* and *Pe* intensities were used to measure the emission anisotropy of the sample.

For FRAP experiments, the sample was imaged in TIRF illumination mode with a single camera. During photobleaching step, the field diaphragm was manually closed to spatially restricting the illumination to the centre of the imaging field. Before and after photobleaching, the aperture was dilated to its fully opened position to record the fluorescent signal. Images were acquired at regular intervals (2-5 s) to record the whole recovery phase. All the experiments were run with Micro-manager software (micro-manager.org). Images were saved as 16-bit (EMCCD) or 12-bit (sCMOS) TIFF files.

**TIRF FRAP analysis**

TIRF FRAP was used to assess the integrity of fluorescent SLBs and to semi-quantitatively estimate the F-actin affinity of the membrane-actin linkers. Time-lapse movies were recorded encompassing pre-beach fluorescence intensity and the post-bleach fluorescence recovery. Images were processed in Fiji. After background subtraction with buffer only images, mean intensity values were measured from the bleached spot (ROI size was kept same as the size of the bleach spot on an immobile sample, ~11 µm in this case) and a reference region (drawn away from the bleach spot). For photobleach correction, a normalized reference profile was obtained by dividing the intensity values from the reference region by its mean ‘pre-bleach’ intensity (mean of the reference region in the frames recorded before photobleaching). Next, the intensity values from the bleached spot were divided by the normalized reference values to obtain photobleach corrected values. Finally, a normalized recovery curve was obtained by dividing all the photobleach-corrected values by the mean ‘pre-bleach’ value (mean of the photobleach-corrected values from the pre-bleach frames). For effective radius (Re) estimation, a line scan of the bleached spot was taken from the first image after photobleaching as well as from the average projection of the pre-bleach frames. The profiles from the bleached spot were then divided by the respective values from the pre-bleach profiles to obtain a normalized bleach profile. The normalized bleach profile and normalized recovery profiles were analysed suing Matlab.

Re was calculated by fitting the normalized bleach profile to the following equation. It accounts for diffusion of the fluorophore during the photobleaching step).

$$f(r) = \exp\,(-K \exp\left(-\frac{2(x-g)^2}{R_e^2}\right) \qquad (1)$$

K informs about the extent of photobleaching, x is the distance in microns, g is the peak position of the inverse Gaussian function, and $R_e$ is the effective radius in microns (Kang et al., 2012).

To calculate the diffusion parameters, the normalized recovery profile was fitted to the following function:

$$f(\mathrm{t}) = \left\{ 1 - \frac{\mathrm{K}}{1 + \frac{R_n^2}{R_e^2} + \frac{8Dt}{R_e^2}} \right\} * \mathrm{M_F} + (1 - \mathrm{M_F})\mathrm{F_0} \qquad (2)$$

The lateral diffusion coefficient (D) is obtained from the fit. $F_0$ is the normalized bleach depth, $F_\infty$ is the saturated fluorescence recovery, D is the lateral diffusion coefficient of the fluorophore in $\mu m^2 s^{-1}$, t is the time in s, MF is the mobile fraction, Rn is the nominal or actual radius of the bleach spot (I estimated Rn by photobleaching a non-diffusing bilayer; one can also use a fluorophore-covered coverslip). K, and Re are used from the bleach profile, as discussed before.

Alternatively, one can use equations 6-8 (discussed in section 2.10.1.6) or fit the recovery profile to an exponential model $A_1(1 - e^{-t/\tau_1})$ to determine the half-time of recovery, $\tau_{1/2}$, from which the D can be calculated by using the following equation:

$$D = \frac{\gamma_d . \omega^2}{4\tau_{1/2}} \qquad (3)$$

$\gamma_d$ is a constant that depends upon the shape of the illumination beam and the extent of photobleaching and is usually fixed at 0.88 (for K < 3), $\omega$ is the radius of the photobleached spot, and $\tau_{1/2}$ is the time at which the fluorescence recovery is 50% (Axelrod et al., 1976).

**TIRF FRAP to quantify lateral diffusion of SLB-bound binder proteins**

TIRF FRAP was used to quantify lateral mobility of Utr-ABD, Ezr-ABD and Ezr-KA on the supported lipid bilayer (SLB) in the before and after adding F-actin. 10nM of each binder was added to the bilayer after which the FRAP measurements of the free binder were performed. After the measurement 250nM of Actin was added (500nM for HSE-KA) incubated for 40 mins after which the bilayer was washed and the FRAP measurements of binder that is binding to actin were performed. Photobleaching experiments were done on Nikon-TiE Epi Fluorescence microscope with an attached TIRF arm. One pre-bleach images was recorded before a small round spot was bleached with 100% laser power (by closing the field aperture). The photobleaching time was empirically adjusted to get about 30-35% photobleaching. The exact size of the bleach spot (Rn) was empirically determined by photobleaching a fixed sample. Images were recorded every 6 seconds and each trace was taken for 50 frames (total 5 mins). Three to five trace was recorded per SLB; 10 to15 traces were collected for each binder.

Using Matlab and Fiji, the acquired image stacks were converted into normalized fluorescence recovery profiles. Image stacks were first background-subtracted in Fiji. To correct for photobleaching and other global intensity fluctuations, mean intensity of each photobleached region was divided by the mean intensity of its reference. The reference region was drawn outside of the photobleached region, at least as large as the bleached region, and far enough from the bleached region to avoid any influence of the initial photobleaching. Finally, each trace was normalized by dividing all the photobleach-corrected values by mean prebleach values.

Next, we determined the mean effective bleaching radius of each iteration of the bleaching from the postbleach fluorescence profiles. The postbleach fluorescence profile of the spot was normalized by their respective prebleach profile. The normalized postbleach profiles were fitted to the following equation:

$$F(x) = e^{-K.e^{\frac{-2(x-g)^2}{Re^2}}}$$

where, K informs about the extent of photobleaching, x is the distance in microns, g is the peak position of the inverse Gaussian function, and $R_e$ is the effective radius in microns (Kang et al., 2012). $R_e$ was determined from the fit for each trace. ($R_e$ compensates for any lateral diffusion during the photobleaching step, which varies for the binders based on their diffusion coefficient). Similarly, we determined the actual radius of the bleached region ($R_n$) by photobleaching a fixed sample using the same settings. $R_e$ and $R_n$ were used as initial conditions for fitting the recovery time profile to the FRAP model described by Kang et al. (Kang et al., 2012) and the Diffusion Coefficient and the Mobile Fraction were estimated as the fitted parameters.

**Image analysis for *in-vitro* experiments**

Multicolour TIRF images of fluorescent F-actin, membrane-actin linkers and sMM-II were processed in Fiji software. First, the backgrounds were subtracted with buffer-only reference images. Next, flat-field correction was done on all the images in each channel: First, we created an average projection of multiple “pre-myosin” images from different regions of the bilayer (“pre-myosin” images were captured in each channel before myosin was added to the system). Next, a Gaussian blurred map ($\sigma$ = 50 pixels for EMCCD images; 80 pixels for sCMOS images) was created from the Average projection image. The map was converted to a 32-bit image, and all the pixels were then divided by the mean value of the whole image to obtain a normalized correction map. Normalized maps were thus generated for each channel. All the images were then divided by their respective normalized maps to correct for non-uniform illumination.

When necessary, correction of small interference fringes (flat-filed correction works for only large-scale patterns) in actin-binder channels was done using a similar method as flat-field correction. First, a minimum projection of the “pre-myosin” images was created for each channel, followed by applying a Gaussian blur filter ($\sigma$ = 3 pixels) on the Minimum projection image, followed by conversion to 32-bit and mean-normalization. All the images in each binder channel were then divided by their respective maps.

Next, photobleaching correction of time-lapse images was done using ‘Photobleach Correction’ plugin in Fiji. Depending on the temporal intensity decay profile of the whole image, an ‘Exponential’ or ‘Simple ratio’ method was used. Each stack was corrected separately and then the three channels were merged into a multichannel Hyperstack. Next, using Hyperstack-Reg (or MultiStackReg) plugin in Fiji, the Hyperstack was corrected for any xy translational movement by ‘Translation’ or ‘Rigid Body’ transformation methods. Finally, the dark pixels along the boundary were cropped, the aligned Hyperstack was split into single channels and saved as 16-bit TIFF stacks for further analysis in Matlab.

**Aster segmentation**

For intensity quantifications and analysis, actin channel was used for segmentation to demarcate ‘asters’ from the background region (see Fig. S3e). First, specific frames in actin channel were selected based on distinctness of actin structures. More than one aster reference images (aster cycles) were selected from image stacks with multiple actomyosin contraction cycles at different time points. Using intensity and size-based thresholding in Matlab, each selected aster reference image was segmented into “asters” (i.e. the regions within distinct actin clusters) and the “background” region (i.e. the remaining region outside of the asters) (Fig. S3f). If the aster density was too high or the aster shape was too complicated, circular ROIs were manually drawn in Fiji. The ROI were then converted into a binary mask, and the coordinates of each segmented object were then exported to Matlab for further processing.

**2D surface plots (intensity color maps)**

To make 2D surface plots from multicolour time-lapse image stacks, aster segmentation was performed as described earlier (Fig. S3f). Next, the centroid of each selected aster was located— we used intensity-weighted centroids of solid asters or geometric centroids of ring-like asters. Each aster region was then demarcated into concentric, non-overlapping circles or annuli of increasing radii, starting from the centroid and spanning the whole aster. Mean intensity (integrated intensity divided by the no. of pixels in the circle) was calculated for each circle of each aster at all the time points. The radial intensities thus obtained from each aster were then normalized by dividing the values by their respective mean pre-myosin intensity values (average intensity for each circle from the images recorded before myosin was added to the system. Typically, >10 'pre-myosin' images were recorded). To compare radial profiles between asters of different radii, normalized radial distance was calculated for each aster by dividing the absolute radii (of all the circles in an aster) by the radius of the largest ring (radius of the whole aster). The coordinates of asters from the actin channel were then exported to the binder channels to determine the normalized radial profiles of the binders. Normalized radial profiles for all the time points were then plotted as a 2D surface plot with x = normalized radius, y = time in s, and z (color map) = mean normalized intensity (fold change with respect to pre-myosin intensity level) of all the asters from one experiment. To estimate the extent of binder sorting in the asters under different phases, we also calculated the normalized binder ratio ($I_A/I_B$) by dividing the pre-bleach normalized radial profile of the stronger binder channel (A) by that of the weaker binder channel (B) for each time point. The binder ratio was plotted as a separate surface plot of the normalized binder ratio (e.g., Fig. S4a-b and j-k).

**Time-dependent mean intensity plots**

To make time dependent mean intensity plots from multicolour time-lapse image stacks, the same user defined segmentation was used to demarcate asters from the background. Mean intensity was calculated for each aster (whole aster region with no radially resolved information) at each time points in all three channels. The intensities were normalized by dividing the intensity values from each aster by its mean 'pre-myosin' intensity values. Normalized radial distance for each aster was calculated by dividing the absolute radii of all the circles by the radius of the aster. Normalized radial profiles (fold change with respect to the pre-myosin level) for each aster was then plotted as a line plot with x = time in seconds, and y = mean normalized intensity of all the asters from one experiment. Data from the three channels was represented in three different colors; this color coding is consistently maintained throughout the paper. For each profile, bold line represents the mean and spline error bar represents (± SEM) of data from all the analysed asters. We also calculated the mean binder ratio ($I_A/I_B$) by dividing the pre-myosin normalized radial profile of the stronger binder channel (A) by that of the weaker binder channel (B) for each time point. The binder ratio was plotted separately as mean (bold line) ± SEM (spline error bar) of data from all the analysed asters from one experiment against time (s) (Fig. S3h).

**Radial intensity plots**

To plot normalized radial intensity profiles from multiple steady-state aster images (recorded as snapshots around the peak intensity of actin in the asters), each image in actin channel was segmented into "asters" and "background" as discussed earlier. Similarly, each aster was demarcated into concentric, non-overlapping circles, starting from the centroid and spanning the whole aster. Mean intensity was calculated from each circle of each aster. Intensity values thus obtained from all the asters in each image were then normalized by dividing the intensity values by mean intensity value of the whole image (typically ~50 μm x ~50 μm or bigger). Normalized radial distance (from the centroid) of each aster was calculated by dividing the absolute radii (of all its circles) by the radius of the aster. Mean normalized intensity (solid line) ± standard error of mean (spline error bar)) of multiple asters from multiple repeats of the experiment was then plotted against the normalized radial distance. The segmentation information was imported to the two binder channels to calculate the radial intensity profiles. Data from each channel was plotted in a different color. These plots were used to classify the sorting patterns of the binders.

***Segregation parameter analysis of in vitro data***

To quantify microphase segregation of actin and actin-binding proteins in vitro, we computed a spatially resolved segregation parameter (ϕ) from time-lapse TIRF microscopy data. Image stacks for binder 1 and binder 2 were background-subtracted, flat-field corrected and corrected for any XY drift and photobleaching in Fiji, as described earlier. The images were then imported as 16-bit TIFF files to MATLAB and converted to double precision data. In each channel, the intensity was normalized to 'pre-myosin' intensity levels. A temporal rolling average (5-15 frame window) was applied to reduce noise and enhance detection of transient clustering. For each time point, local segregation parameter (ϕ) was calculated within a sliding 3×3-pixel spatial window as ϕ = IA - IB/IA + IB), where $I_A$ and $I_B$ are the normalized intensities of A (stronger) and B (weaker) binder within the local spatiotemporal window. ϕ maps were generated for each time point and exported as 32-bit TIFF stacks, using a fixed color scale for cross-frame comparison. In this context, ϕ = 0 indicates complete mixing, ϕ = +1 o r-1 indicates complete phase separation by complete exclusion of B, or A, respective. Intermediate values reflect partial segregation.

#### Figure legends for Supplementary figures:

**Figure S1: In vitro reconstitution and two different activity states of contractile asters.** **a**, Schematic of the in-vitro reconstitution system, assembled hierarchically from a glass-supported lipid bilayer, an ABP linker protein, capped actin filaments and myosin-II minifilaments. The system was imaged with a multicolour TIRF microscope at 100-150x magnification. **b**, Snapshot from TIRF microscopy showing a region of the bilayer with polar actin asters at the steady state (before arresting in the ATP-depleted, jammed state) and the locally clustered membrane-bound linker protein. Scale bar = 5 μm. **c-e**, 2D intensity colour maps showing the radial intensity distribution of F-actin-Atto635, HYE (YFP-tagged linker) and myosin II-Atto565 in the remodelling aster state. The intensity values in each channel were normalized to the pre-myosin levels. Y-axis represents the normalized distance from the aster centroid, x-axis shows time (s). **f**, Mean intensity (± SEM) of actin (black), myosin (brown) and HYE (magenta) during the remodelling aster phase (n = 14). **g-i**, 2D intensity colour maps showing normalized radial intensity of F-actin, HYE and myosin II in the non-remodelling aster phase. **j**, Mean intensity (± SEM) from 14 individual asters of actin (black), myosin (brown) and HYE (magenta) in the jammed aster phase. **k**, Dissolution of jammed asters (connected aster phase (Köster et al., 2016)) following addition of ATP (1 mM). Notable enhancement of colocalization between F-actin and HYE can be observed 4 s post ATP addition. Scale bar = 10 μm.

**Figure S2: Different assays to measure F-actin affinity of the binders. a**, Principle of ABP-F-actin co-sedimentation assay. Binder 1 with strong affinity for F-actin co-sediments more with F-actin than Binder 2, which has a weaker actin affinity. **b**, Calibration curve converting background-subtracted band integrated densities into protein amounts. **c**, Plot showing actin-normalized (μmol/μmol) and free fractions of the ABP, fitted to the Hill-Langmuir function to determine $K_D$ (the equilibrium dissociation constant or $k_{off}/k_{on}$), $B_{max}$ (maximum binding capacity) and n (Hill coefficient). **d**, Principle of the single-molecule dwell-time assay using TIRF microscopy. **e**, Snapshot showing F-actin-Alexa488 immobilized on the surface via Biotin-Streptavidin interactions (as shown in d). **f**, Snapshot shows fluorescent actin-binding molecules (Atto-565) dwelling on F-actin in the evanescence field. **g**, Kymograph showing single binding events; the length of each line reflects the dwell time ($1/k_{off}$) of each binding event. **h-j**, Results from fitting the co-sedimentation data to the Hill-Langmuir model to obtain $K_D$, $B_{max}$, and n for Utr-wt (h), Ezr-wt (i), Ezr-KA (j). Estimated values are shown in each plot, along with gel images from SDS-PAGE of the pelleted fractions. Fitting was performed to minimize RMSE, and values > 0.1 were excluded from fitting (shown as red crosses). **k-n**, Data from single molecule assay showing frequency plots of F-actin dwell times for the three binders: Utr-wt (k), Ezr-wt (l), and Ezr-KA (m); (n) is the control showing non-specific binding of Ezr-wt to a surface passivated with BSA/BSA-Biotin without F-actin. Data in k-m were fitted to a multi-exponential model, one for the non-specific binding component and the second one for the characteristic F-actin dwell time. Utr-wt is the strongest binder, followed by Ezr-wt and Ezr-KA. The affinity values from both the assay were used to obtain the duty ratios, $k_{on}/(k_{on} + k_{off})$. **o**, Normalized FRAP (fluorescent recovery after photobleaching) profiles of the three binders in the absence and presence of F-actin on an SLB. **p**, Lateral diffusion coefficient (D) of the binders in the absence and presence of F-actin. D was quantified by fitting the fluorescence recovery profile to f(t)= {1- K/(1+(R_n^2)/(R_e^2)+8Dt/(R_e^2))} *M_F+(F_∞-M_F ) F_0 as described in (Kang et al., 2012). **q**, Ratio of D in the presence of F-actin (D) to that in its absence of F-actin ($D_0$). Utr-wt exhibits the lowest D/Do ratio, indicating its highest affinity for F-actin among the three actin binders.

**Fig. S3**: **Image processing and data analysis.** **a-c**, TIRF images of F-actin-Atto633 (grey), Ezr-wt (YFP tagged) (magenta, stronger F-actin binder) and Ezr-KA (SNAP549) (green, weaker F-actin binder). **a**, Uniform distribution of the adapter proteins on the lipid bilayer with no F-actin. **b**, Shows the steady state distribution of the adapters in the presence F-actin before myosin addition. Ezr-wt colocalized with F-actin. **c**, Snapshot of actin asters in the dynamic steady state, showing actin enrichment in dense actomyosin asters with local clustering of Ezr-wt and exclusion of Ezr-KA. **d**, Actin asters in the ATP-depleted, rigor state, where both binders loose spatial patterning due to lack of actin dynamics. **e-i**, Image processing and segmentation. **e**, Time-lapse image stacks are background-subtracted, flat-field corrected and loaded to Matlab for segmentation. **f**, For segmentation, an image is chosen depending on the distinctness of actin structures in actin channel (Channel 1) during high-ATP or low-ATP state. A threshold is applied based on intensity and size of the detected objects, and the image is segmented into "asters" and the "background" region. The aster mask is manually filtered to remove artefacts. Pixel indices and centroid position of each accepted aster region are saved and applied to other two channels for further analysis. **g**, Surface plots of normalized radial intensity profiles, where x= normalized distance from the aster centroid, y = time (s), z = intensity fold-change with respect to pre-myosin levels. **h**, 2D normalized intensity plots showing mean intensity values from multiple aster regions (whole aster regions without radial information) over time. Data from the three channels is plotted in three different colours: consistently maintained throughout this paper. Bold lines represent the mean and shaded error bars represent ± SEM. **i**, Normalized radial intensity profiles of asters from multiple experiments, where x = normalized radial distance and y = fold-change with respect to pre-myosin intensity levels. Solid line represents the mean; the spline error bar represents ± SEM. These plots are used to classify molecular patterns of the actin binders.

**Figure S4**: **Active clustering and spatial sorting of actin-binding proteins (ABPs) driven by remodelling contractile asters. a-b**, Binder sorting ratio (BSR), defined as the ratio of normalized intensities of the stronger to weaker binder (Ezr-wt/Ezr-KA) in the remodelling (a) and jammed aster state (b). BSR values of 1, > 1, or < 1 indicate no net enrichment, enrichment, or depletion of the stronger binders with respect to the weaker binder, respectively. **c-d**, Time course of normalized mean intensities of actin, Ezr-wt, Ezr-KA and BSR in the entire aster regions [n = 21 for remodelling state (c); n = 40 for jammed state (d)]. Bold lines denote mean; shaded error bar represents SEM. **e**, Bar plot comparing aster formation rate between remodelling (50 asters) and jammed states (41 asters). Each data point denotes the rate values obtained from fitting each individual actin intensity trace. **f**, Scatter plot showing the correlation between actin growth rate (rate of actin enrichment within aster regions) and BSR growth rate (rate at which BSR increases within asters). **g**, Scatter plot showing correlation between actin growth rate and BSR amplitude (extent of binder sorting). h, Time course of normalized actin intensity (SEM in the shaded region) from two remodelling aster phases, showing the aster lifetime of each aster cycle and the remodelling time between the two. All the traces in each aster cycle were aligned by peak timing after which a gaussian fit was applied to the mean trace to measure aster lifetime. **i**, summary table of dynamic parameters quantified for Ezr-wt: Ezr-KA (1:1 density ratio) condition in remodelling and jammed aster states, derived from mean intensity profiles aligned by peak timing (note: intensity profiles in a-g and j-p are not peak aligned). The aster lifetime was quantified using the full width half maximum (FWHM) of a Gaussian fit, while rate constants were measured by exponential fitting in Matlab. Error ranges represent the 95% fitting confidence interval. Remodelling time was calculated over two remodelling aster cycles, centred at 474 s and 840 s, respectively, before the system entered the jammed aster phase. **j-k**, BSR (Utr-wt/Ezr-wt) measured in remodelling (j) and jammed states (k). **l-m**, Normalized mean intensities of actin, Utr-wt, Ezr-wt and BSR in remodelling (l) and jammed states (m). Bold lines denote mean; shaded error bar represents SEM. **n**, Bar plot comparing aster formation rate in remodelling and jammed states (17 asters each). Each data points in the plot denotes the rate value obtained from fitting each individual actin intensity trace. **o**, Scatter plot showing correlation between actin growth rate and BSR growth rate. **p**, Scatter plot showing correlation between the actin growth rate and BSR amplitude (the extent of binder sorting).

**Figure S5**: **Increased relative density reduces exclusion of the weak binder from asters. a-b,** Binder sorting ratio (BSR; Utr-wt/Ezr-wt) measured in remodelling (a) and jammed states (b) at 1:4 density ratio. **c-d**, Time course of normalized mean intensities of actin, Utr-wt, Ezr-wt and BSR in remodelling (c) and jammed states (d). Bold lines denote mean; shaded error bar represents SEM. **e-h,** Normalized radial intensity colormaps (**e-h**) display spatial distribution of F-actin (e), Utr-wt (f) and Ezr-wt (g), and BSR (h) following ATP re-addition to jammed asters. **i**, Time course of normalized mean intensities is shown in (i).

**Figure S6**: **Actin binders with equal actin affinity co-cluster in both remodelling and jammed states. a-e**, Spatial organization during the remodelling aster state (n = 13 asters). TIRF images of a representative remodelling aster (**a**). **b-d**, Normalized radial intensity colormaps of F-actin, Ezr-wt1 (YFP-tagged) and Ezr-wt2 (SNAP549 labelled) show strong transient mixed clustering of the two binders with equal actin binding affinities. Both binders exhibit strong, transient mixed clustering in the remodelling state. **e,** Radial map of ϕ shows no increase during peak actin enrichment (~110 s), despite a >2.5 peak fold change in F-actin, suggesting robust mixed coclustering. **f-j**, Spatial organization during the jammed state (n = 21 asters). **f**, TIRF images of a representative jammed state aster. **g-i**, Normalized radial intensity colormaps of F-actin, Ezr-wt1 and Ezr-wt2 show localized mixed clustering (t ~180 s) with a slow decay profile. **j**, Radial map of ϕ confirms persistent mixed clustering of Ezr-wt1 and Ezr-wt2 with no significant change in the order parameter throughout the clustering phase.

**Figure S7**. **Active retrograde flows at membrane protrusions drive sorting of membrane proteins with differential actin affinities**. **a-c,** TIRF images shows GFP-nanobody-Alexa546 and FR-Fab-Star635 fluorescence in representative cells for the three binder conditions: Ezr-RA: Ezr-RA (non-binder control, a), Ezr-wt: Ezr-wt (equal binder control, b) and Utr-wt: Ezr-wt (differential binder test, c). Corresponding FR/GFP ratio colormaps are shown in the main figure (Fig. 7c). **d-f**, Representative kymograph close ups display GFP and FR intensities, highlighting protrusion and retraction events. Corresponding FR/GFP ratio profiles are shown in Fig. 7e. **g-h**, Radial profiles of normalized FR (solid lines) and GFP (dashed lines) intensities along protrusions (g) and retractions (h) for each case (see figure legends). **i**, Boxplots (with data overlap) showing the distribution of edge velocities across the three conditions (N= 3 independent experiments; ns = non-significant by two sample t-test). **j-k**, scatter plots showing the correlation between FR/GFP ratio and retraction rate for Ezr-RA: Ezr-RA (j), Ezr-wt: Ezr-wt (k) and Utr-wt: Ezr-wt (k) pairs. No clear correlation is observed in any of the three conditions.

#### Supplementary Movie Legends:

**Supplementary Movie 1 (Fig2, remodeling state)** – Representative time-lapse sequences (montages described in Fig 2b) showing TIRF images of F-actin (grey, left panel), Ezr-wt (magenta, middle panel) and Ezr-KA (green, right panel) during the high-ATP remodeling aster state. The movie starts at the pre-myosin state, where both the binders are more uniformly distributed, and following myosin-II addition (not shown), the asters are formed, and the two binders are segregated by the dynamic asters. Images were acquired at a15-second interval. The scale bar is 3 µm. The displayed movie segment covers 765 s and plays at 10 frames per second.

**Supplementary Movie 2 (Fig2, jammed state)** – Representative time-lapse sequences (montages described in Fig 2g) showing TIRF images of F-actin (grey, left panel), Ezr-wt (magenta, middle panel) and Ezr-KA (green, right panel) during the low-ATP jamming aster state. The movie starts at the pre-myosin state, the jammed aster begins around frame 44 in the movie, gradually driving the two binders to a mild segregated state before actin arrests in a static rigor state. Images were acquired every 15 seconds. The scale bar is 3 µm. The displayed movie segment covers 1590 s and plays at 15 frames per second.

**Supplementary Movie 3 (Fig3, aster simulation)** – Representative time-lapse sequences (density profiles described in Fig 3a), showing images from Monte Carlo simulation of molecules A (red) and B (green), illustrating their spatial distribution before, during, and after an aster appears in the 2d simulation field. The aster is turned on at t = 50000, and turned off at t = 55000 simulation steps, during which A is clustered and B is excluded from the aster region. After the aster disappears, the density levels return to the baseline.

**Supplementary Movie 4 (Fig4, remodeling state)** – Representative time-lapse sequences (montages described in Fig 4a) showing TIRF images of F-actin (grey, left panel), Utr-wt (yellow, middle panel) and Ezr-wt (magenta, right panel) during the remodeling aster state. Notice how the relatively stronger binder, Utr-wt, exhibits a robust enrichment with actin and the relatively weaker binder, Ezr-wt, exhibits a modest enrichment in the asters. Images were acquired every 10 seconds. The scale bar is 3 µm. The displayed movie segment covers 700 s and plays at 20 frames per second.

**Supplementary Movie 5 (Fig4, jammed state)** – Representative time-lapse sequences (montages described in Fig 4f) showing TIRF images of F-actin (grey, left panel), Utr-wt (yellow, middle panel) and Ezr-wt (magenta, right panel) during the remodeling aster state. Notice how the relatively stronger binder, Utr-wt, exhibits a robust enrichment with actin. The relatively weaker binder, Ezr-wt, exhibits mixed patterning followed by a transient bull's eye like pattern before it is eventually excluded from the jammed asters. Images were acquired every 10 seconds. The scale bar is 3 µm. The displayed movie segment covers 910 s and plays at 30 frames per second.

**Supplementary Movie 6 (Fig6, spreading cell)** – Representative time-lapse sequences (described in Fig 6) showing TIRF images of mouse embryonic fibroblasts spreading on fibronectin-coated glass. The cells co-expressing FR-TM-Ezr-wt (red, top left panel) and GFP-TM-Utr (green, top right panel) were fluorescently labelled with either anti-FR Fab (MoV19Fab-Star635) or anti-GFP nanobody (GFP-nanobody-Alexa546). The lower, left panel shows the Ezr-wt/Utr-wt ratio over time, and the lower, left panel outlines the cell shape over time. The Ezr-wt/Utr-wt ratio is elevated at the cell edge during lamellipodial membrane protrusion and decreases towards the cell center, indicating sorting along the centripetal direction of the actin flow. Images were acquired every 5 seconds. The scale bar is 5 μm.

Fig. S1

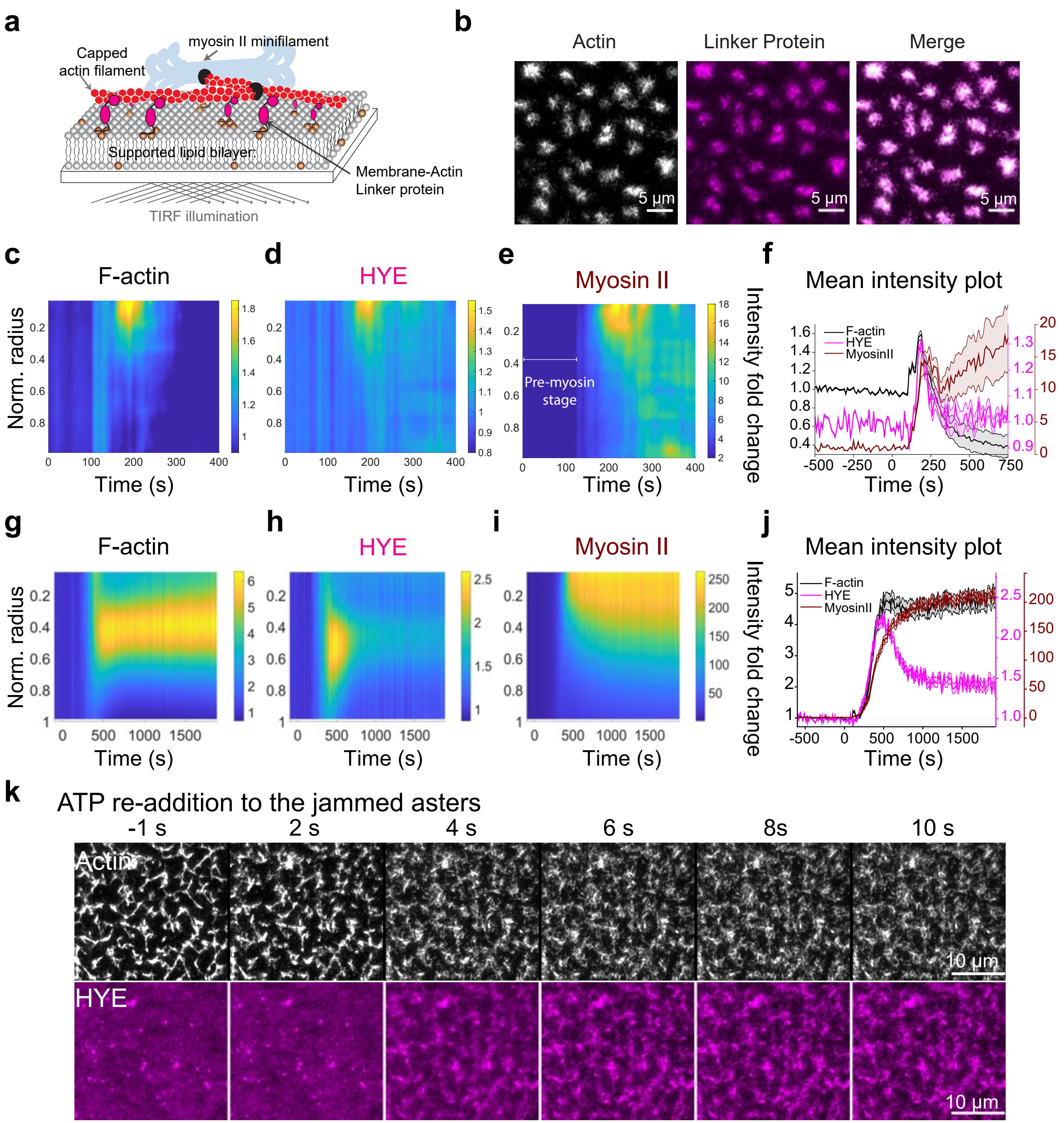
a
myosin II minifilament
Capped actin filament
Supported lipid bilayer
Membrane-Actin Linker protein
TIRF illumination
b
Actin
Linker Protein
Merge
5 µm
c
F-actin
d
HYE
e
Myosin II
Pre-myosin stage
f
Mean intensity plot
F-actin
HYE
MyosinII
Norm. radius
Time (s)
Intensity fold change
g
F-actin
h
HYE
i
Myosin II
j
Mean intensity plot
k
ATP re-addition to the jammed asters
-1 s
2 s
4 s
6 s
8s
10 s
Actin
HYE
10 µm

Fig. S2

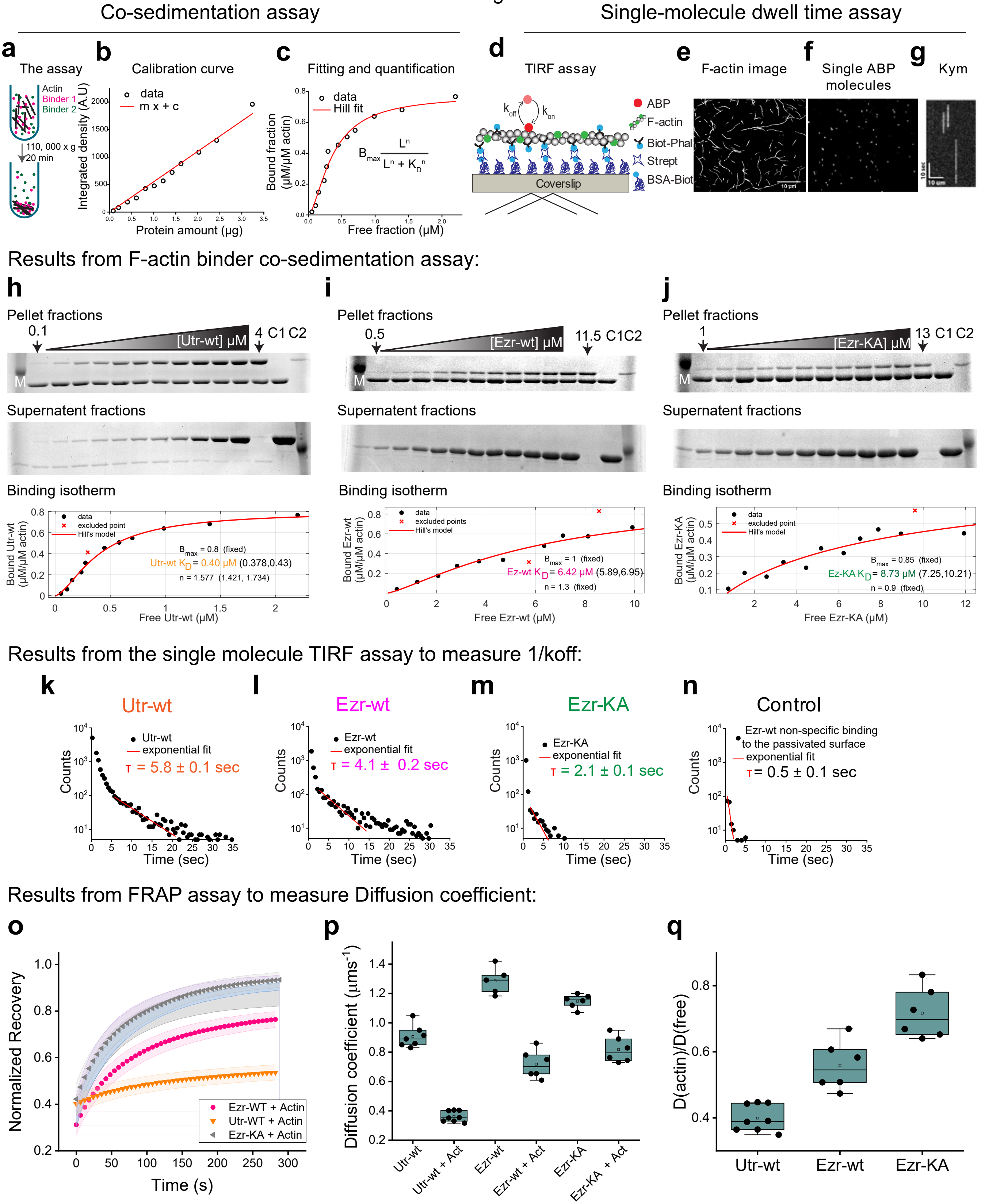

Co-sedimentation assay
Single-molecule dwell time assay
a The assay
Actin
Binder 1
Binder 2
110, 000 x g
20 min
b Calibration curve
data
m x + c
Integrated density (A.U)
Protein amount (µg)
c Fitting and quantification
data
Hill fit
Bound fraction (µM/µM actin)
Free fraction (µM)
d TIRF assay
k_off
k_on
ABP
F-actin
Biot-Phal
Strept
BSA-Biot
Coverslip
e F-actin image
f Single ABP molecules
g Kym
Results from F-actin binder co-sedimentation assay:
h
Pellet fractions
0.1
[Utr-wt] µM
4 C1 C2
M
Supernatent fractions
Binding isotherm
data
excluded point
Hill's model
B_max = 0.8 (fixed)
Utr-wt K_D = 0.40 µM (0.378,0.43)
n = 1.577 (1.421, 1.734)
Bound Utr-wt (µM/µM actin)
Free Utr-wt (µM)
i
Pellet fractions
0.5
[Ezr-wt] µM
11.5 C1C2
M
Supernatent fractions
Binding isotherm
data
excluded points
Hill's model
B_max = 1 (fixed)
Ez-wt K_D = 6.42 µM (5.89,6.95)
n = 1.3 (fixed)
Bound Ezr-wt (µM/µM actin)
Free Ezr-wt (µM)
j
Pellet fractions
1
[Ezr-KA] µM
13 C1 C2
M
Supernatent fractions
Binding isotherm
data
excluded point
Hill's model
B_max = 0.85 (fixed)
Ez-KA K_D = 8.73 µM (7.25,10.21)
n = 0.9 (fixed)
Bound Ezr-KA (µM/µM actin)
Free Ezr-KA (µM)
Results from the single molecule TIRF assay to measure 1/koff:
k Utr-wt
Utr-wt
exponential fit
τ = 5.8 ± 0.1 sec
l Ezr-wt
Ezr-wt
exponential fit
τ = 4.1 ± 0.2 sec
m Ezr-KA
Ezr-KA
exponential fit
τ = 2.1 ± 0.1 sec
n Control
Ezr-wt non-specific binding to the passivated surface
exponential fit
τ = 0.5 ± 0.1 sec
Counts
Time (sec)
Results from FRAP assay to measure Diffusion coefficient:
o
Normalized Recovery
Time (s)
Ezr-WT + Actin
Utr-WT + Actin
Ezr-KA + Actin
p
Diffusion coefficient (µms^-1)
Utr-wt
Utr-wt + Act
Ezr-wt
Ezr-wt + Act
Ezr-KA
Ezr-KA + Act
q
D(actin)/D(free)
Utr-wt
Ezr-wt
Ezr-KA

Fig. S3

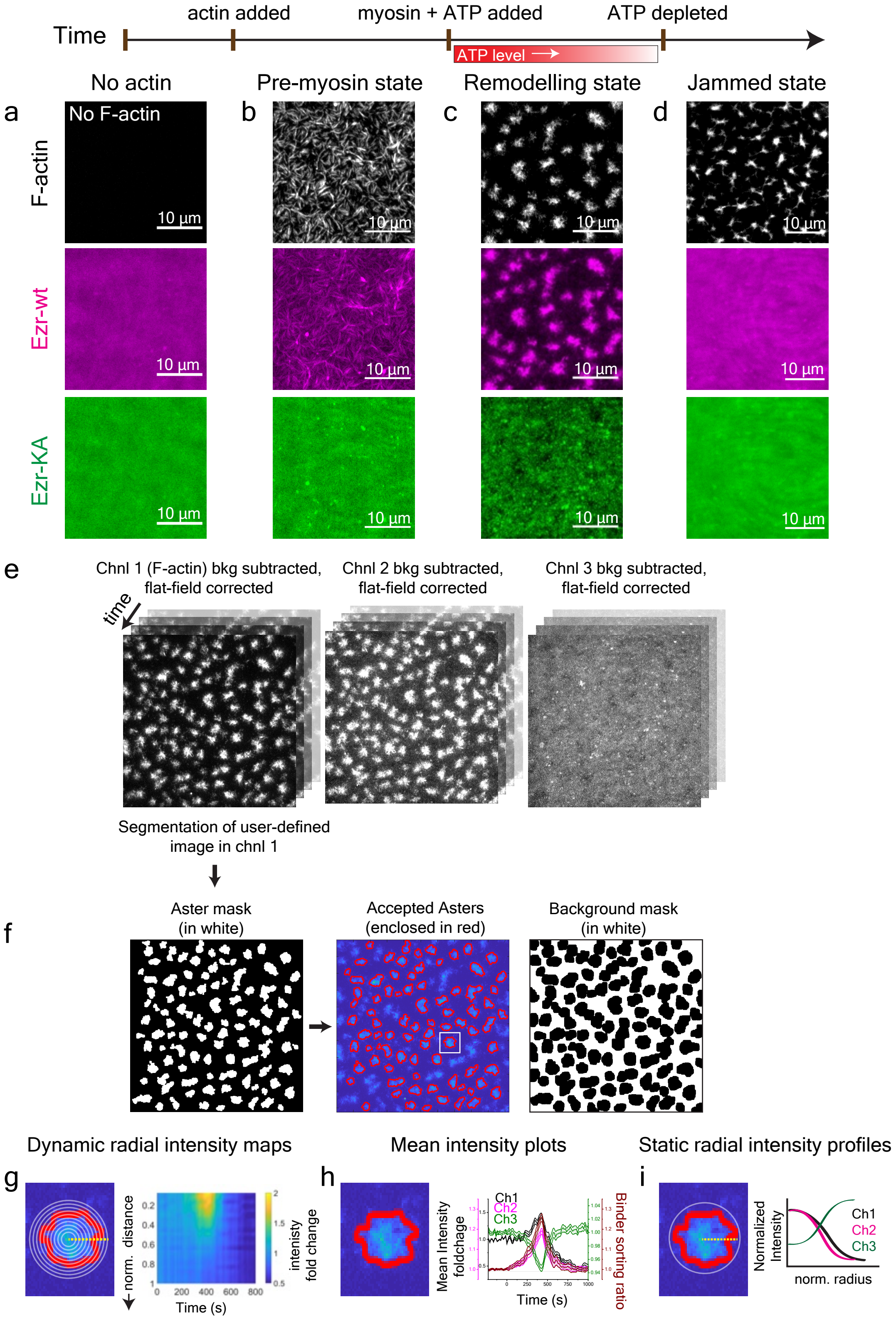

actin added
myosin + ATP added
ATP depleted
Time
ATP level
No actin
Pre-myosin state
Remodelling state
Jammed state
a
b
c
d
F-actin
Ezr-wt
Ezr-KA
No F-actin
10 µm
e
Chnl 1 (F-actin) bkg subtracted, flat-field corrected
Chnl 2 bkg subtracted, flat-field corrected
Chnl 3 bkg subtracted, flat-field corrected
time
Segmentation of user-defined image in chnl 1
f
Aster mask (in white)
Accepted Asters (enclosed in red)
Background mask (in white)
Dynamic radial intensity maps
Mean intensity plots
Static radial intensity profiles
g
h
i
norm. distance
intenisty fold change
Time (s)
Mean Intensity foldchage
Binder sorting ratio
Ch1
Ch2
Ch3
Normalized Intensity
norm. radius

### Fig. S4

Ezr-wt : Ezr-KA (1:1 molar density)

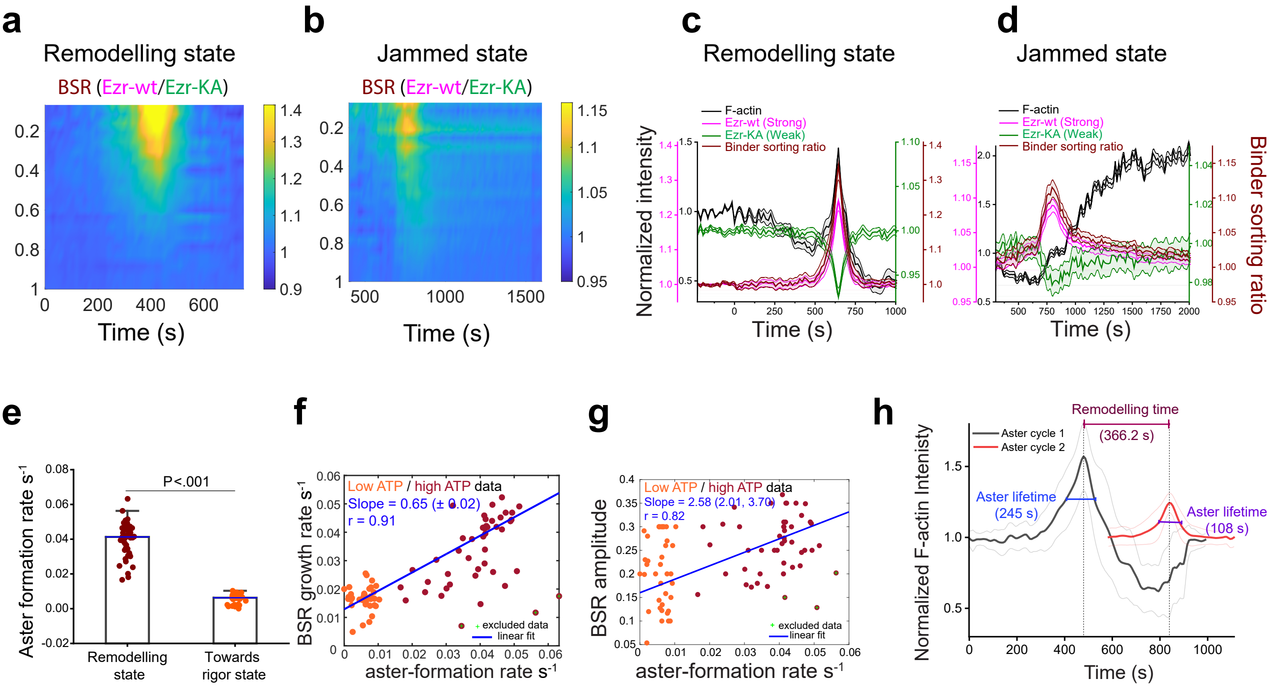


i Dynamic parameters quantified for Ezr-wt : Ezr-KA (1:1 molar density) case (B1 here is Ezr-wt)

| Parameter | Remodelling state | Jammed state |
|---|---|---|
| Actin aster lifetime (s) | 245 (±89) | 1033 (±4.8) |
| B1 cluster lifetime (s) | 205 (±21) | 148.3 (±46.5) |
| Actin aster rise rate (s^-1) | 0.0083 (±0.0013) | 0.0043 (±0.0015) |
| B1 clustering rate (s^-1) | 0.0062 (±0.0014) | 0.0159 (±0.0038) |
| Actin aster decay rate (s^-1) | (-) 0.0053 (±0.0023) | NA |
| B1 cluster decay rate (s^-1) | (-) 0.0149 (±0.0024) | (-) 0.0068 (±0.0013) |
| Remodelling time (s) | 366.2 (±17.92) | NA |
| Remodelling rate (s^-1) | 0.00273 (±0.00013) | NA |

Utr-wt : Ezr-wt (1:1 molar density)

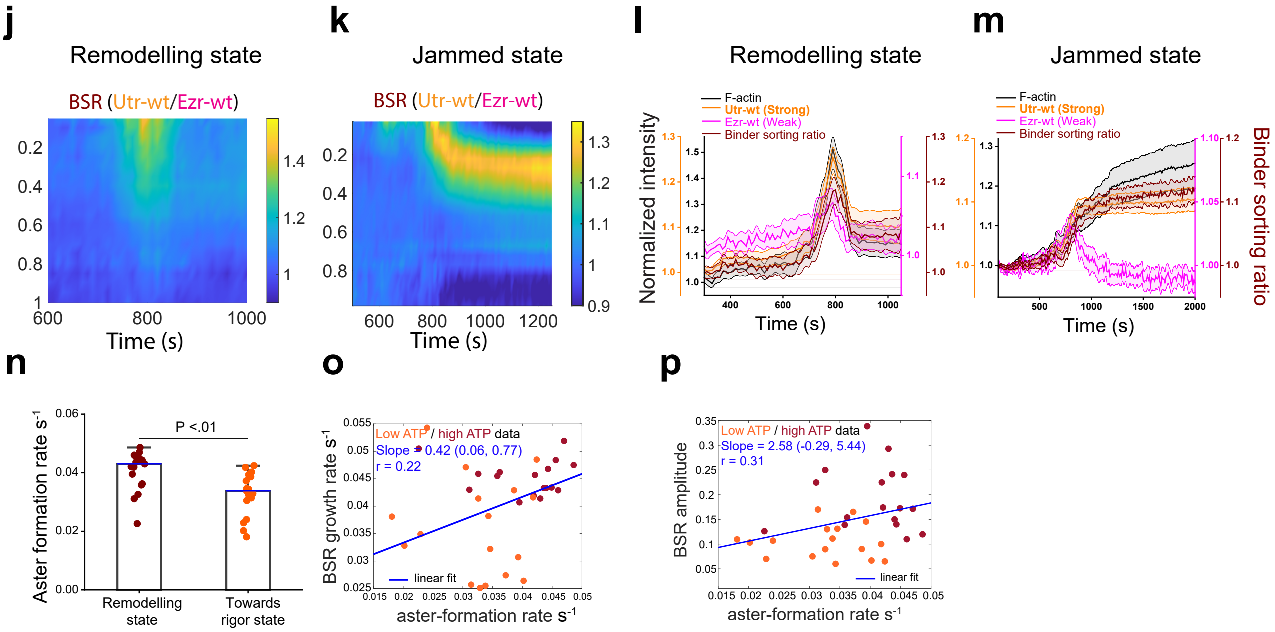

Fig. S5

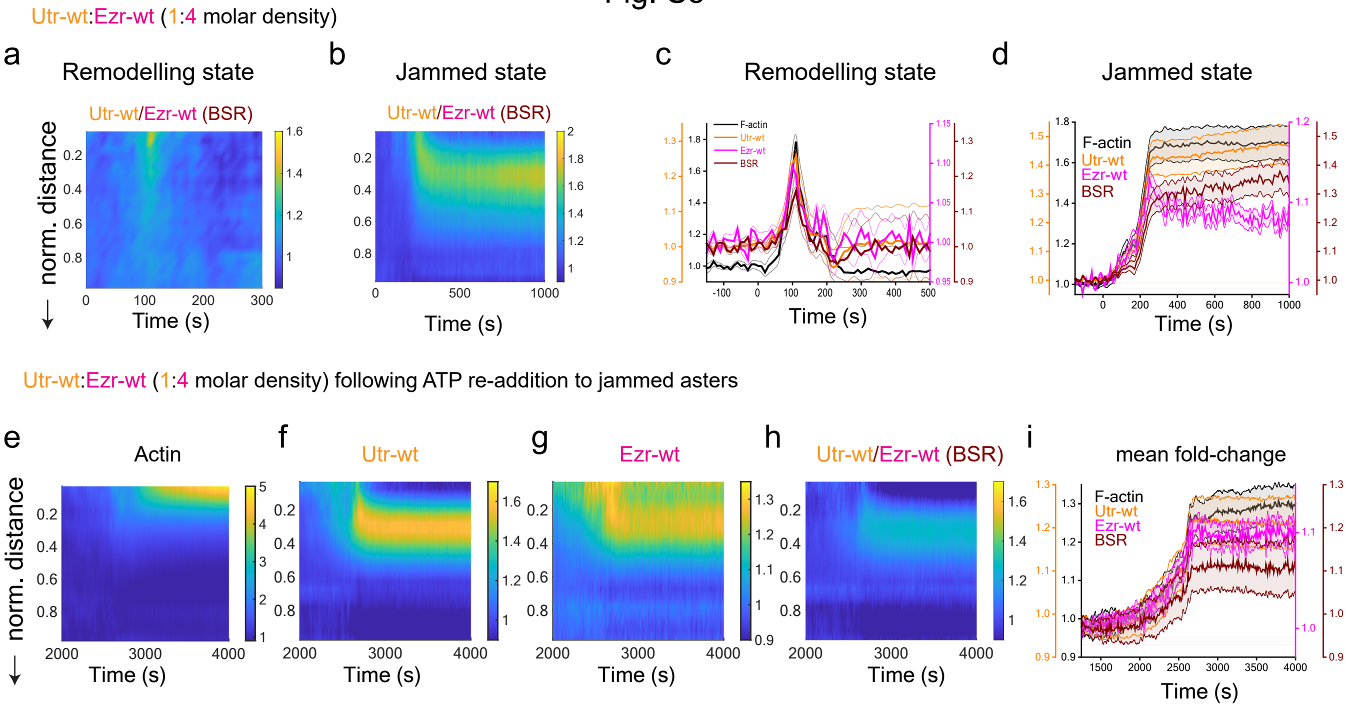

Utr-wt:Ezr-wt (1:4 molar density)
a
Remodelling state
Utr-wt/Ezr-wt (BSR)
norm. distance
Time (s)
b
Jammed state
Utr-wt/Ezr-wt (BSR)
Time (s)
c
Remodelling state
F-actin
Utr-wt
Ezr-wt
BSR
Time (s)
d
Jammed state
F-actin
Utr-wt
Ezr-wt
BSR
Time (s)
Utr-wt:Ezr-wt (1:4 molar density) following ATP re-addition to jammed asters
e
Actin
norm. distance
Time (s)
f
Utr-wt
Time (s)
g
Ezr-wt
Time (s)
h
Utr-wt/Ezr-wt (BSR)
Time (s)
i
mean fold-change
F-actin
Utr-wt
Ezr-wt
BSR
Time (s)

### Fig. S6

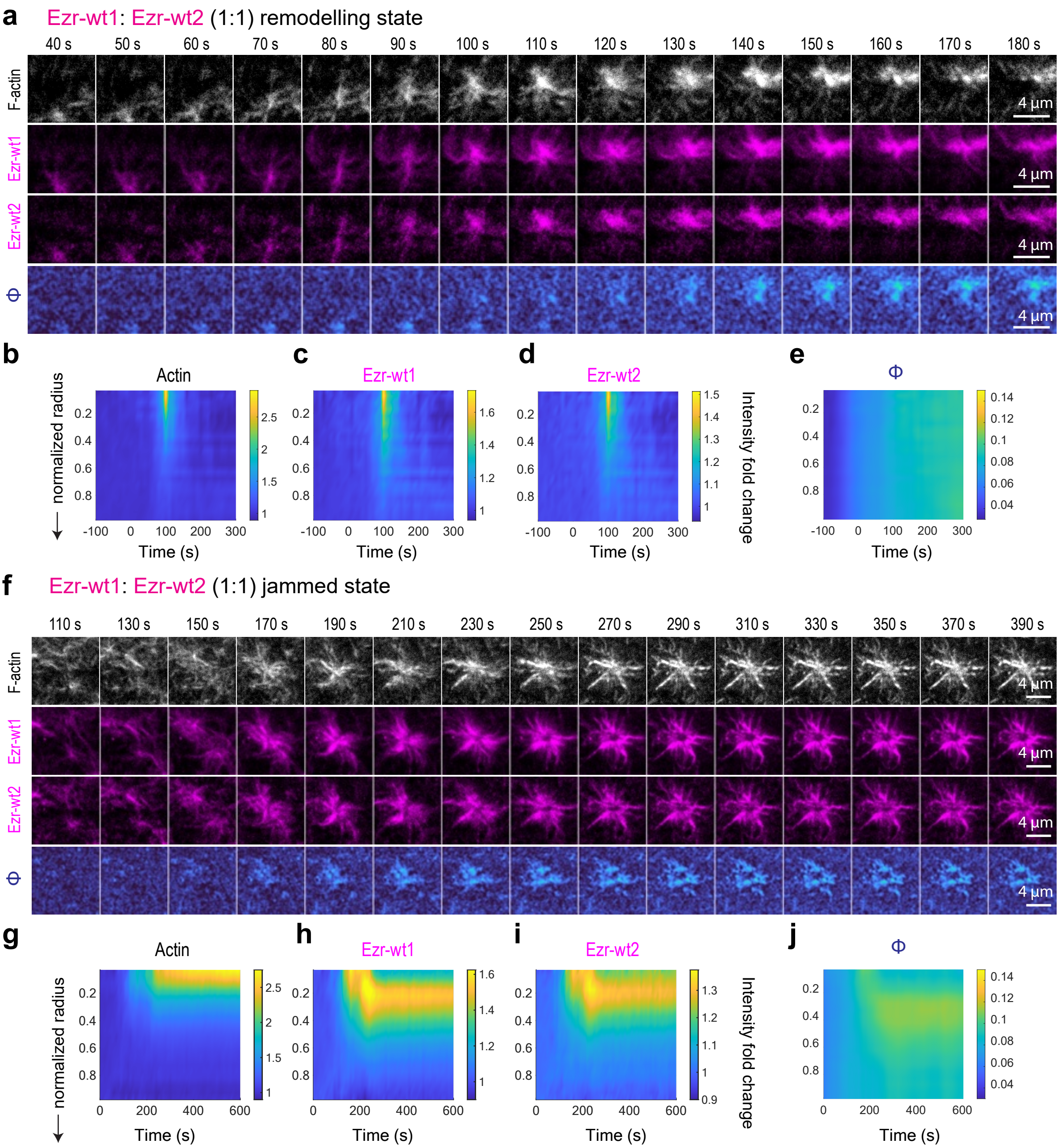
a
Ezr-wt1: Ezr-wt2 (1:1) remodelling state
40 s
50 s
60 s
70 s
80 s
90 s
100 s
110 s
120 s
130 s
140 s
150 s
160 s
170 s
180 s
F-actin
Ezr-wt1
Ezr-wt2
Φ
4 µm
b
Actin
c
Ezr-wt1
d
Ezr-wt2
e
Φ
normalized radius
Time (s)
Intensity fold change
f
Ezr-wt1: Ezr-wt2 (1:1) jammed state
110 s
130 s
150 s
170 s
190 s
210 s
230 s
250 s
270 s
290 s
310 s
330 s
350 s
370 s
390 s
F-actin
Ezr-wt1
Ezr-wt2
Φ
4 µm
g
Actin
h
Ezr-wt1
i
Ezr-wt2
j
Φ
normalized radius
Time (s)
Intensity fold change

### Fig. S7

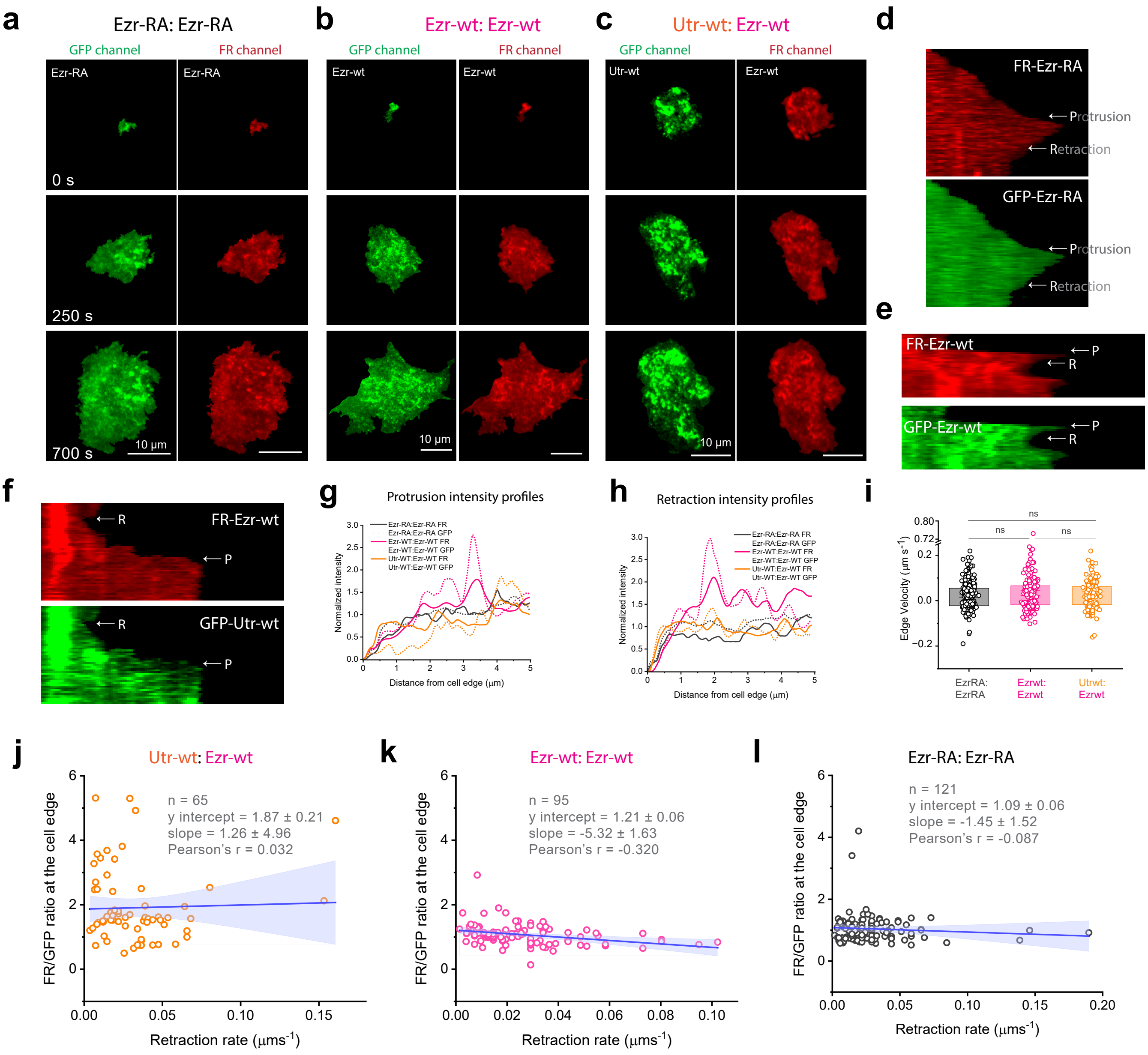
a
Ezr-RA: Ezr-RA
GFP channel
FR channel
Ezr-RA
Ezr-RA
0 s
250 s
700 s
10 µm
b
Ezr-wt: Ezr-wt
GFP channel
FR channel
Ezr-wt
Ezr-wt
10 µm
c
Utr-wt: Ezr-wt
GFP channel
FR channel
Utr-wt
Ezr-wt
10 µm
d
FR-Ezr-RA
Protrusion
Retraction
GFP-Ezr-RA
Protrusion
Retraction
e
FR-Ezr-wt
P
R
GFP-Ezr-wt
P
R
f
FR-Ezr-wt
R
P
GFP-Utr-wt
R
P
g
Protrusion intensity profiles
Ezr-RA:Ezr-RA FR
Ezr-RA:Ezr-RA GFP
Ezr-WT:Ezr-WT FR
Ezr-WT:Ezr-WT GFP
Utr-WT:Ezr-WT FR
Utr-WT:Ezr-WT GFP
Normalized intensity
Distance from cell edge (µm)
h
Retraction intensity profiles
Ezr-RA:Ezr-RA FR
Ezr-RA:Ezr-RA GFP
Ezr-WT:Ezr-WT FR
Ezr-WT:Ezr-WT GFP
Utr-WT:Ezr-WT FR
Utr-WT:Ezr-WT GFP
Normalized intensity
Distance from cell edge (µm)
i
ns
ns
ns
Edge Velocity (µm s$^{-1}$)
EzrRA: EzrRA
Ezrwt: Ezrwt
Utrwt: Ezrwt
j
Utr-wt: Ezr-wt
n = 65
y intercept = 1.87 ± 0.21
slope = 1.26 ± 4.96
Pearson's r = 0.032
FR/GFP ratio at the cell edge
Retraction rate (µms$^{-1}$)
k
Ezr-wt: Ezr-wt
n = 95
y intercept = 1.21 ± 0.06
slope = -5.32 ± 1.63
Pearson's r = -0.320
FR/GFP ratio at the cell edge
Retraction rate (µms$^{-1}$)
l
Ezr-RA: Ezr-RA
n = 121
y intercept = 1.09 ± 0.06
slope = -1.45 ± 1.52
Pearson's r = -0.087
FR/GFP ratio at the cell edge
Retraction rate (µms$^{-1}$)
