## Supplementary Information-III for "Active flows drive clustering and sorting of membrane components with differential affinity to dynamic actin cytoskeleton"

### Supplementary Information for
### Active flows drive clustering and sorting of membrane components with differential affinity

Bhat et al

#### I. KINETIC MONTE CARLO SIMULATION MODEL OF PROTEINS WITH DIFFERENTIAL BINDING AFFINITIES BY LOCALISED ACTIVE CONTRACTILE STRESSES

Here we describe a coarse-grained off-lattice model for the dynamic patterning of proteins with differential binding affinities driven by localised active contractile stresses generated by a active fluid in 2d. This is motivated by our earlier models described in Refs. [1–3].

##### A. Multi-component supported bilayer membrane

We begin with a minimal description of the lateral dynamics of two cell surface proteins, A and B, in the *in vitro* supported bilayer. Both these proteins bind and unbind to the underlying reconstituted actomysoin cortex with fixed affinities which are different for protein A and protein B. The molecules of each of the proteins are assumed to have very weak homophilic interactions. However, they form transient clusters in response to their interactions with the actomysoin cortex which drives these molecules via active contractile stresses flucuating in space and time.

In our coarse grained description we represent all protein molecules as spheres, that reside on the bilayer membrane treated as a flat surface of lateral dimension $L \times L$ and area $A = L^2$. The total number of molecules in the plane is fixed at $N$, constituted by $N_A$ molecules of protein A and $N_B$ molecules of protein B. Thus, total number density on the plane always remain fixed.

The molecules of the two protein species interact via a modified Lenard-Jones potential (MLJP) of the form,

$$\begin{aligned} V(X_i^{\alpha}, X_j^{\beta}) &= 4\left[\varepsilon^r_{\alpha\beta}\left(\frac{\sigma_{\alpha\beta}}{r_{ij}^{\alpha\beta}}\right)^{12} - \varepsilon^a_{\alpha\beta}\left(\frac{\sigma_{\alpha\beta}}{r_{ij}^{\alpha\beta}}\right)^{6}\right], \text{ if } \alpha = \beta \\ &= 4\varepsilon^r_{\alpha\beta}\left(\frac{\sigma_{\alpha\beta}}{r_{ij}^{\alpha\beta}}\right)^{12}, \text{ if } \alpha \neq \beta \end{aligned} \tag{1}$$

Eq 1 describes the repulsive as well as attractive interaction between $i$-th molecule of component $\alpha$ and the $j$-th molecule of component $\beta$. Here we consider short-range repulsive interactions among all molecules, irrespective of species identity whose strength is given by $\varepsilon^r_{\alpha\beta}$. We consider longer-range attractions only between molecules of same type, give by $\varepsilon^a_{\alpha\beta}$. Both interaction strengths are expressed in units of thermal energy $k_BT$ ($k_B$ is the Boltzmann constant and $T$ is the absolute temperature). The value of $\varepsilon^r_{\alpha\beta}$ is taken to be a constant ($= k_BT$) for all combinations of $\alpha$ and $\beta$ in our simulations. However, we tune $\varepsilon^a_{\alpha\alpha}$ and $\varepsilon^a_{\beta\beta}$ to assess the interplay of clustering due to strength of homophilic attractions and active driving. Furthermore, $\sigma_{\alpha\beta}$ represents the hard-core length scale and $r_{ij}^{\alpha\beta} = |X_i^{\alpha} - X_j^{\beta}|$ is the distance between the two interacting molecules. For simplicity, we take $\sigma_{\alpha\beta} = \sigma$ for all $\alpha, \beta$.

##### B. Active contractile stresses from actomyosin cortex

The cortical actomyosin reconstituted on top of the supported bilayer applies stochastic contractile stresses on the membrane proteins when they bind to it. Historically, the dynamic contractile stresses are realized through a Poisson distributed birth-death process that is exponentially correlated in time, with a characteristic timescale $\tau_{rem}$ [1–4] which is similar to the remodeling timescale observed in the experiments. Moreover, these contractile stresses are strongly correlated over a spatial scale $\xi$ and uncorrelated otherwise. The amplitude of such stresses are finite and contractile over certain regions in space. We designate one such region as:$\{\Omega_{\mathbf{x}_a}\}$ with radius $\xi$ centred around $\mathbf{x}_a$, $a = 1, ...., n$, given by,

$$\begin{aligned} \Sigma(x,t) &= -\Sigma_0\, \hat{\mathbf{r}}_a, \text{ if } x \in \Omega_{\mathbf{x}_a} \\ &= 0, \text{ if } x \notin \Omega_{\mathbf{x}_a} \end{aligned} \tag{2}$$

where $\hat{\mathbf{r}}_a$ is the unit radial vector from the centre of the domain $\Omega_{\mathbf{x}_a}$ and $\Sigma_0$ is the amplitude of the stress. We create an ensemble of $n$ such active stress events $n$, meaning the coverage of nonzero active stress is given by $S_{active} = n\pi\xi^2/L^2$. In our current framework, we consider only one such active stress event ($n = 1$) at any point of time during a given simulation.

#### II. DYNAMICS OF ACTIVE SEGREGATION REALIZED BY OFF-LATTICE KINETIC MONTE-CARLO METHOD

The dynamics of the cell surface proteins, subject to both equilibrium and active forces, are described in terms of a Master equation for the time evolution of the probability distribution, $P(\{X_i^\alpha\},\{\Omega_{\mathbf{x}_a}\},t)$. We solve the Master equation using a kinetic Monte Carlo approach, where we specify the updates for the positions $\{X_i^\alpha\}$ of the protein molecules and placement $\{\Omega_{\mathbf{x}_a}\}$ of the active stress events.

The configuration in the membrane surface is updated by a sequence of moves determined both by equilibrium pair-potentials (Eq. 1) and by forces arising from the local contractile stress if the particles are in the region $\{\Omega_{\mathbf{x}_a}\}$ (Eq. 2). The details of the transition rates associated with these moves are described below and in Fig. 1b. Note that one time step of this kinetic Monte-Carlo scheme, $\Delta t$ is given by $N$ attempts of position updates some fraction of which are diffusive moves governed by equilibrium forces and the rest are active advection moves driven by contractile stresses. All the position updates conform to Periodic boundary conditions in both X and Y directions.

##### A. Position updates governed by equilibrium pair-potentials

Choosing any protein molecule of species $A$ or $B$ with uniform probability, we attempt a position update by a small random displacement $\delta r_d$, also uniformly distributed, with the following probability,

$$\begin{aligned} p_{eq} &= 1, \quad \text{if } \Delta\mathcal{E} \le 0 \\ &= e^{-\Delta\mathcal{E}/k_BT}, \quad \text{if } \Delta\mathcal{E} > 0 \end{aligned} \tag{3}$$

where $\Delta\mathcal{E}$ is the difference in energy before and after the update, and is obtained from the inter-particle potentials described by Eq. 1. One such move is illustrated in Fig. S1A. This is the implementation of usual Monte-Carlo update of molecular systems akin to the Metropolis algorithm and obeys detailed balance [5, 6].

##### B. Position updates governed by contractile stresses

This applies only to molecules that are within the contractile regions $\Omega_{\mathbf{x}_a}$ and bound to it with affinities, $K_A$ or $K_B$, according to the species chosen being A or B, respectively. They move preferentially towards the centre $\mathbf{x}_a$ with a radial velocity, $v_a$. In our simulations, $v_a$ is implemented by a series of attempted radial displacements each of magnitude $\delta r_a$ with a fixed advection probability $\Sigma_0$. Such moves, illustrated in Fig. S1B, often incur high energy costs and thus violate detailed balance. To stabilize the simulations, we disallow such moves when the resultant energy cost is higher than $20k_BT$, the approximate amount of energy released upon hydrolysis of a single molecule of ATP.

##### C. Adjustment of displacements

The initial magnitude of $\delta r_d$ ($\sim 0.2\sigma$) and $\delta r_a$ ($\sim 0.05\sigma$) are taken to be only a few percentage of the average linear size of the protein molecules. These values, however, are adjusted regularly during the simulation with a fixed frequency in order to ensure an overall acceptance of 40% position updates which is a standard protocol for Monte-Carlo simulations [5, 6].

##### D. Ratio of equilibrium and active moves

In an off-lattice setting, the fraction of diffusion and advective moves is determined by the Peclet number which is essentially the ratio of advective to diffusive motion. In our simulations, every protein molecule outside the range of the region $\Omega$ undergoes diffusion and within $\Omega$ undergoes advection towards the centre of the region. For a diffusion coefficient, $D$ and an advection speed $v_a$, we can define a Peclet number $P_e = v_a\sigma/D$. We define: $D \approx \sigma^2/(4\tau_d)$ where $\tau_d$ is the diffusion timescale, the duration taken by a diffusing molecule to travel a distance equal to its average diameter $\sigma$. Here we consider diffusion in 2D which is typical of a protein on the plasma membrane. Similarly, we can define an advection timescale $\tau_a$ and write the advection speed as: $v_a = \Sigma_0\sigma/\tau_a$ where the stress amplitude $\Sigma_0$ has been considered as the strength of advection. Here we implement $\Sigma_0$ as the probability of one advection event. Using these two forms, we can derive the following definition of the Peclet number: $P_e = 4\Sigma_0\tau_d/\tau_a$. In our off-lattice setting, we fix the value of $P_e$ and $\Sigma_0$ to estimate the ratio $f = \tau_a/\tau_d = 4\Sigma_0/P_e$. Essentially, $f$ is the ratio of number of diffusive moves ($N_d$) to the number of advective moves ($N_a$) per Monte-Carlo step. In a Monte-Carlo simulation with $N$ molecules, total of $N$ moves, diffusive and advective included, constitute one Monte-Carlo sweep. Therefore, we can determine the number of diffusive moves by $N_d = N\left(\frac{f}{f+1}\right)$ and number of advective moves by $N_a = N - N_d$.

##### E. Choice of Parameters and simulation units

In our simulations we vary the following parameters: (i) Composition of the system, i.e. ratio of $A$ and $B$ molecules, (ii) mean remodeling time of the active stress events $\tau_{rem}$, and (iii) F-actin binding affinities, $K_A$ and $K_B$. We vary the compositions by keeping the total concentration of proteins fixed. We choose protein $A$ to always have a higher actin binding affinity. Thus, to test how composition competes with contractile stresses, we reduce the concentration of $A$ and increase that of $B$. The rest of the parameters of the model, lateral size of the membrane surface $L$, linear scale of the contractile region $\xi$, advection probability $\Sigma_0$, and temperature $T$, are all held fixed. The qualitative results we obtain are independent of these parameters across a large range.

In our simulations, the repulsive interaction strength sets the scale of energy. Most of our simulations are done with $\varepsilon^a_{AA} = \varepsilon^a_{BB} = 0.1 k_B T$, which completely removes the chances of spontaneous segregation. In some specific cases, however, we explicitly change these parameters to study the effect of the attractive interactions on the clustering process. We emphasize that our choice of relative concentration and temperature is not unique and the results hold qualitatively for a wide range of relative concentrations and temperatures.

To express the results of our simulation in physical units, we denote every length scale in our simulation in units of $\sigma = 10\,\text{nm}$, a typical molecular scale. This gives us the spatial scale of contractile stress $\xi = 100\,\text{nm}$ and the lateral size of each bilayer patch $L = 600\,\text{nm}$.

To fix the time unit of our simulations, we measure mean-square displacements of molecules in one of our typical simulations and find that about 10 Monte-Carlo 'timesteps' in our simulations amount to approximately the diffusion timescale of any molecule. Subsequently, we fix the unit of time in our simulations, using a molecular diffusion coefficient of $D \approx 1\ \mu\text{m}^2\ \text{s}^{-1}$ typically observed for cell surface proteins [7]. This gives us each Monte-Carlo timestep: $\Delta t \approx 2.5\ \mu\text{s}$. In these units, a typical simulation run of $10^7$ MC steps corresponds to 25 s. We have also used $\tau_d$ to define remodeling rates for the contractile regions as: $\omega = \tau_d/\tau_{rem}$. So, for $\tau_{rem}$ in range $4-30000$ MC steps, $\omega$ ranges from $3.33 \times 10^{-4}$ to 2.5.

All the simulations with advection are performed with a fixed active radial velocity, implemented as an attempt probability, $\Sigma_0 = 0.8$ and Peclet number, $P_e = 10$. Majority of our simulations with activity are performed with $\tau = 5000$ MC steps $\equiv 12.5$ ms, unless mentioned otherwise.

##### F. Analysis of local density difference

We compute the local density difference order parameter $\phi(r) = \langle n_A(r)\rangle - \langle n_B(r)\rangle$ where $n_A(r)$ and $n_B(r)$ are the radial densities of species A and B, respectively, calculated at a radial distance $r$ from the centre of the contractile domain. We calculate $n_A(r)$ and $n_B(r)$ on a circular grid with spacing $\sigma$ overlaid on the contractile domain. The angular brackets represent averages over time windows for which the contractile domain is present and averages over possible grid points at distance $r$ from the centre of the domain. Such spatiotemporally averaged radial density profiles for A and B are shown in Fig. 3d of the main text. We use these averaged radial densities to compute the density difference, $\phi(r)$.

From $\phi(r)$, we next compute the various Fourier amplitudes. The maximum amplitude, $|\phi(q)|_{max}$ is connected to the wave number $q$ corresponding to the linear size of the largest domain observed in the system [3, 8]. This quantity varies as we observe different types of patterns formed by species A and B within the contractile region. Some of these patterns are illustrated in Fig. S1C. The larger the domain size of any one species within the region, the larger is the value of $|\phi(q)|_{max}$. We define arbitrary cut-offs based on the color-variation on the heatmaps of $|\phi(q)|_{max}$, which help us distinguish among different patterns. The heatmaps exhibit strong dependence on $\tau_{rem}$ of the contractile region. We observe that as $\tau_{rem}$ decreases, the small $|\phi(q)|_{max}$ values which correspond to the mixed phase dominates (Fig. S2).

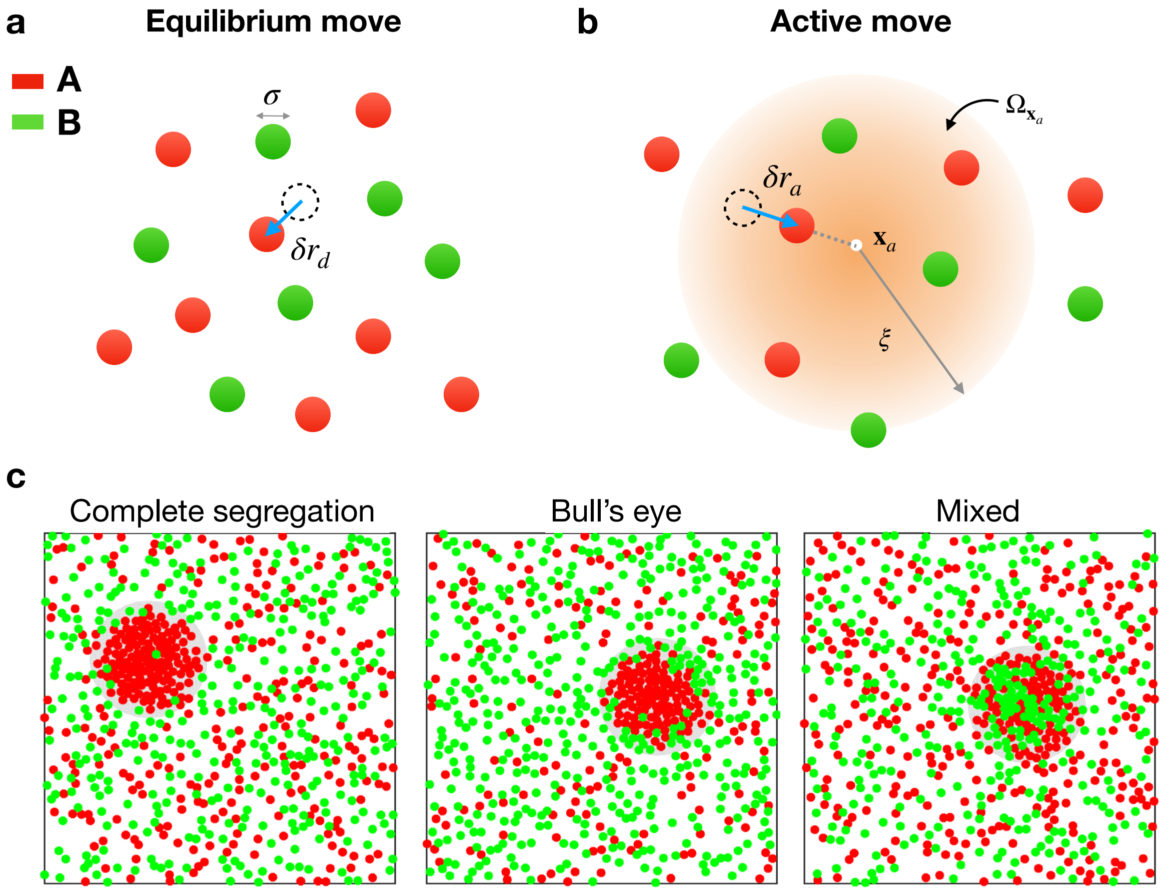


FIG. S1. **Kinetic Monte-Carlo simulation moves: (a) equilibrium moves governed by the pair-potentials as described in Eq. 3. (b) Active moves on protein molecules when they are within the contractile regions $\Omega_{\mathbf{x}_a}$ and bound to it with a rate $K_A$ or $K_B$, respectively. The active moves result in preferentially advecting the molecules towards the centre $\mathrm{x}_a$ with a radial velocity proportional to the contractile stress $\Sigma_0$ (Eq. 2). These moves violate detailed balance. (c) Three different snapshots of A-B patterns formed within the contractile region: left - complete segregation, middle - Bull's eye, and right - mixed.**

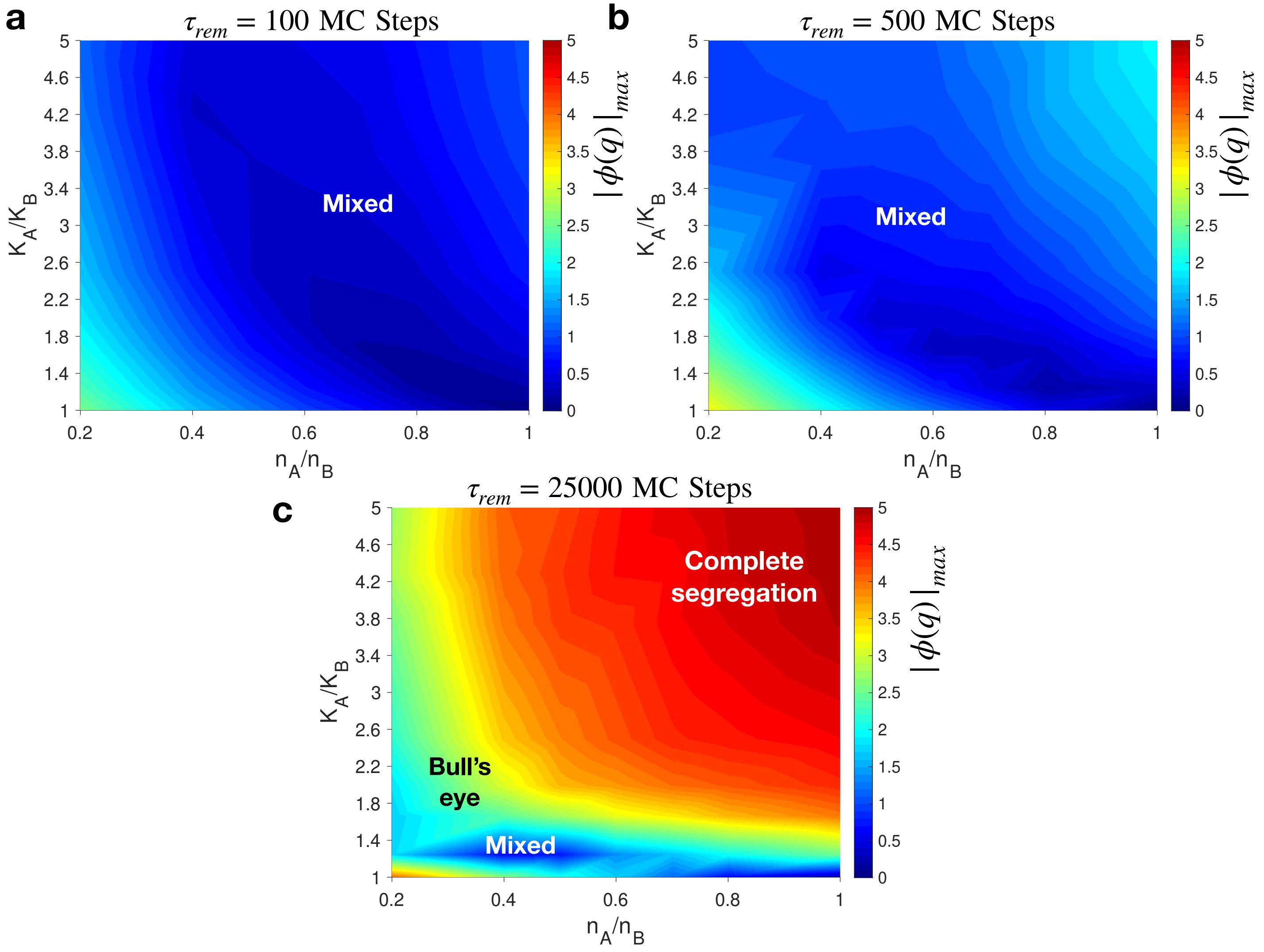


FIG. S2. **Heatmaps of the $|\phi(q)|_{max}$, the maximum Fourier amplitude of local density difference $\phi(r)$ at two different values of remodeling time $\tau_{rem}$ of the contractile region: (a) $\tau = 100$ MC steps, (b) $\tau = 500$ MC steps and (c) $\tau = 25000$ MC steps.**

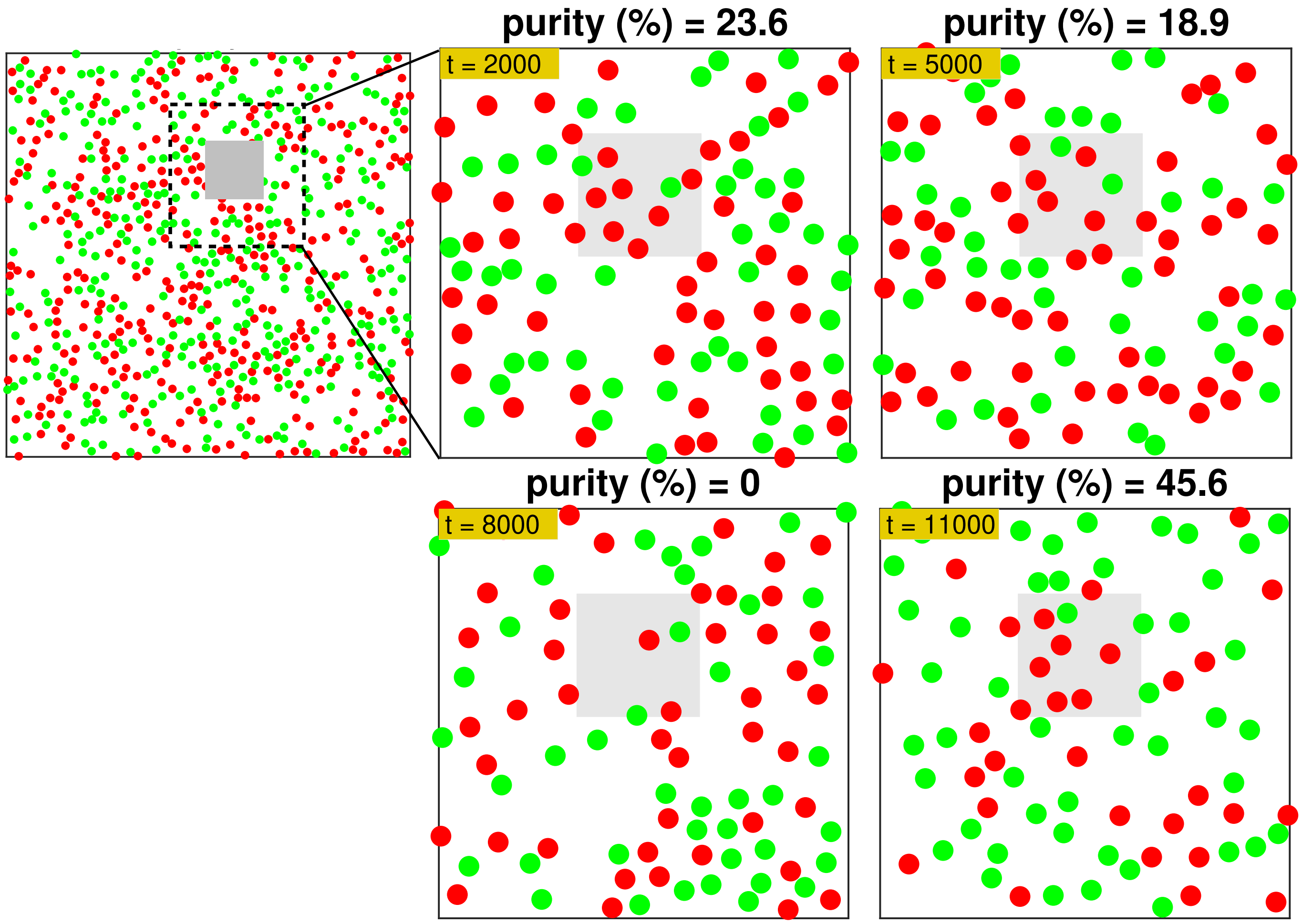


FIG. S3. **Non-equilibrium remodelling can drive chemical purity of clusters. The simulation area is shown on the top left where we have marked a small square region of linear size $6\sigma$ is highlighted. This is the region $\Omega$ where we measure chemical purity $\Delta$ of component A for affinity ratio $K_A/K_B = 1.05$, which means the two species are nearly indistinguishable in terms of affinity to the actomyosin contractile domain. The time evolution of the molecular movements in and out of the gray box is shown in four panels in the right with time-stamps shown on the panels. Here the mean remodeling time of the domain is set at $\tau_{rem} = 10$ MC steps. This corresponds to an extremely high remodeling rate. We compute purity using the following expression: $\Delta = 1 - \langle S_\Omega \rangle / \ln 2$ where $S_\Omega$ is the mixing entropy in the gray region denoted as $\Omega$. $S_\Omega$ is defined as the following: $S_\Omega = -(x_A \ln x_A + x_B \ln x_B$ with $x_A$ and $x_B$ being the number fraction of the two species A and B, respectively (here number fraction and mole fraction is almost identical since the two species have similar molecular weights). When $x_A < x_B$ in $\Omega$, we have set $S_\Omega = \ln 2$, so purity is zero.**
